# Polygenic adaptation from standing genetic variation allows rapid ecotype formation

**DOI:** 10.1101/2021.04.16.440113

**Authors:** Nico Fuhrmann, Celine Prakash, Tobias S. Kaiser

## Abstract

Adaptive ecotype formation is the first step to speciation, but the genetic underpinnings of this process are poorly understood. Marine midges of the genus *Clunio* (Diptera) have recolonized Northern European shore areas after the last glaciation. In response to local tide conditions they have formed different ecotypes with respect to timing of adult emergence, oviposition behavior and larval habitat. Genomic analysis confirms the recent establishment of these ecotypes, reflected in massive haplotype sharing between ecotypes, irrespective of whether there is ongoing gene flow or geographic isolation. QTL mapping and genome screens reveal patterns of polygenic adaptation from standing genetic variation. Ecotype-associated loci prominently include circadian clock genes, as well as genes affecting sensory perception and nervous system development, hinting to a central role of these processes in ecotype formation. Our data show that adaptive ecotype formation can occur rapidly, with ongoing gene flow and largely based on a re-assortment of existing alleles.

## Introduction

The genetic foundations for rapid ecological adaptation and speciation remain poorly understood. Decades of research have focused on identifying single or few major genetic loci in such contexts, with some notable successes ^1–4^. However, many genome-wide association studies have now shown that quantitative trait phenotypes have usually a polygenic basis^5,6^. This implies that also rapid natural adaptation should be expected to occur in a polygenic context^7,8^. Yet, tests of this assumption are still rare, since it is easier to prove the involvement of major locus compared to showing its absence. To show that this occurs in natural populations requires an exceptional biogeographic setting, as well as deep genomic and genetic analysis.

Here we present a study on the postglacial evolution of new ecotypes in marine midges of the genus *Clunio* (Diptera: Chironomidae), which in adaptation to their habitat differ in oviposition behavior and reproductive timing, involving both circadian and circalunar clocks.

Circalunar clocks are biological time-keeping mechanisms that allow organisms to anticipate lunar phase^9^. Their molecular basis is unknown^10^, making identification of adaptive loci for lunar timing both particularly challenging and interesting. In many marine organisms, circalunar clocks synchronize reproduction within a population. In *Clunio marinus* they have additional ecological relevance^11^. Living in the intertidal zone of the European Atlantic coasts, *C. marinus* requires the habitat to be exposed by the tide for successful oviposition. The habitat is maximally exposed during the low waters of spring tide days around full moon and new moon. Adult emergence is restricted to these occasions by a circalunar clock, which tightly regulates development and maturation. Additionally, a circadian clock ensures emergence during only one of the two daily low tides. The adults reproduce immediately after emergence and die few hours later. As tidal regimes vary dramatically along the coastline, populations of this *Atlantic ecotype* of *C. marinus* show local timing adaptations^11–13^ for which the genetic underpinnings are partially known^14,15^.

When *Clunio* expanded into the North after the recess of the ice shield, new ecotypes *evolved* in the Baltic Sea^16–18^ and in the high Arctic^19,20^, which differ from the *Atlantic ecotype* of *C. marinus* in various ways (see Fig. 1 for a summary of defining characteristics of the three ecotypes). In the Baltic Sea the tides are negligible and the *Baltic ecotype* oviposits on the open water, from where the eggs quickly sink to the submerged larval habitat at water depths of up to 20 metres^16,21^. Reproduction of the *Baltic ecotype* happens every day precisely at dusk under control of a circadian clock^22^, without a detectable circalunar rhythm^17^. Near Bergen (Norway) the *Baltic* and *Atlantic ecotypes* were reported to co-occur in sympatry, but in temporal reproductive isolation. The *Baltic ecotype* reproduces at dusk, the *Atlantic ecotype* reproduces during the afternoon low tide^22^. Therefore, the *Baltic ecotype* is currently considered a separate species – *C. balticus*. However, *C. balticus* and *C. marinus* can be successfully interbred in the laboratory^22^.

**Figure 1.**
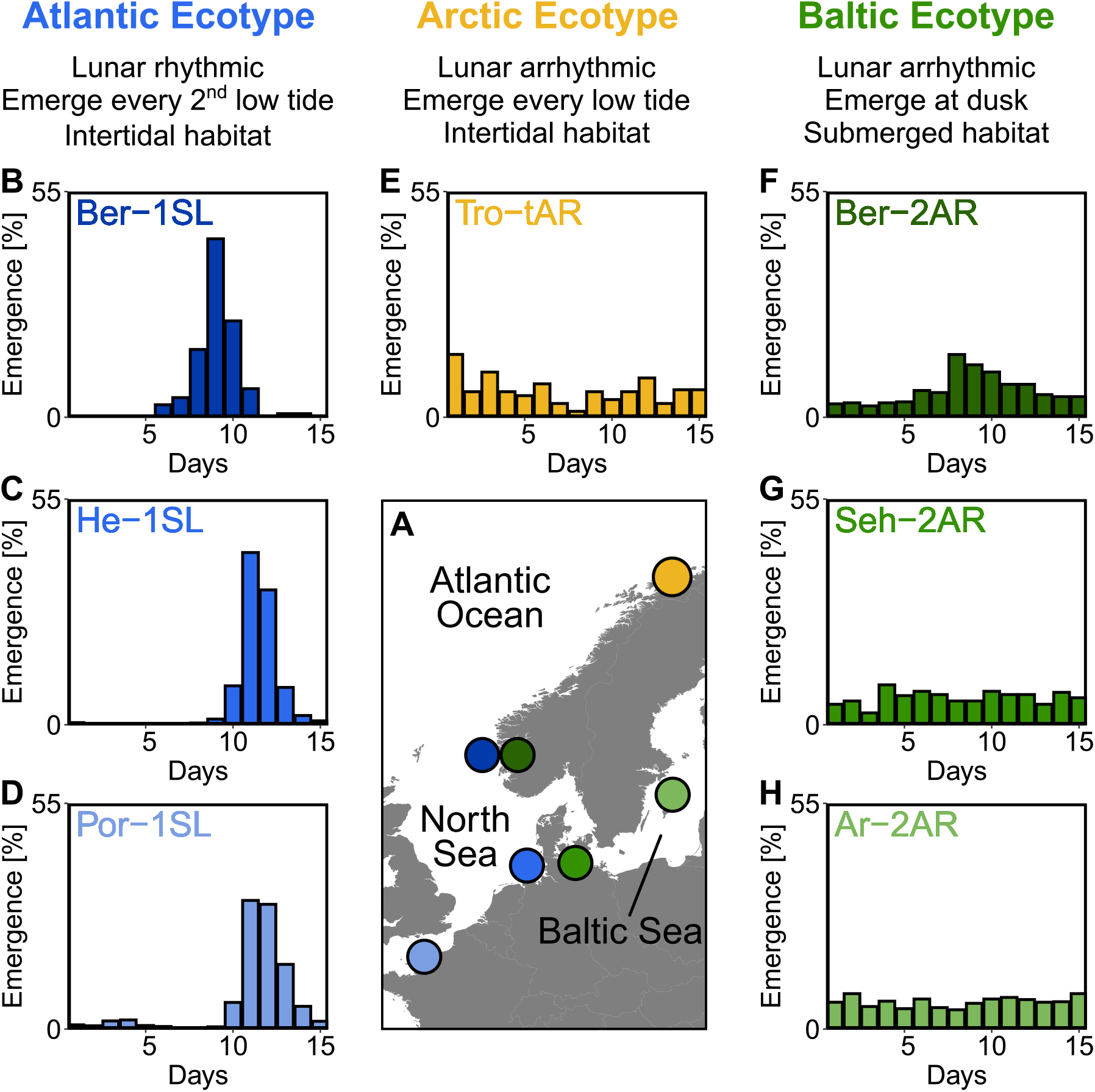
Northern European ecotypes of *Clunio* and their lunar rhythms. The *Atlantic, Arctic* and *Baltic ecotypes* of *Clunio* differ mainly in their lunar rhythms (B-H), circadian rhythms (Suppl. Fig. 1), as well as their habitat and the resulting oviposition behavior (Suppl. Note 1). **(A)** Sampling sites for this study. **(B-H)** Lunar rhythms of adult emergence in corresponding laboratory strains under common garden conditions, with 16 hours of daylight and simulated tidal turbulence cycles to synchronize the lunar rhythm. In *Arctic* and *Baltic ecotypes* the lunar rhythm is absent (E,G,H) or very weak (F). Por-1SL: n=1,263; He-1SL: n=2,075; Ber-1SL: n=230; Tro-tAR: n=209; Ber-2AR: n=399; Seh-2AR: n= 380; Ar-2AR: n=765.

In the high Arctic there are normal tides and the *Arctic ecotype* of *C. marinus* is found in intertidal habitats^19^. During its reproductive season, the permanent light of polar day precludes synchronization of the circadian and circalunar clocks with the environment. Thus, the *Arctic ecotype* relies on a so-called tidal hourglass timer, which allows it to emerge and reproduce during every low tide^20^. It does not show circalunar or circadian rhythms^20^.

The geological history of Northern Europe^23^ implies the Baltic Sea and the high Arctic could only be colonized by *Clunio* after the last glacial ice shield recessed about 10,000 years ago. This inference is also supported by subfossil *Clunio* head capsules in Baltic Sea sediment cores^24^. The new adaptations to the absence of tides in the Baltic Sea and to polar day in the high Arctic must therefore have occurred within this time frame. This makes *Clunio* a particularly interesting model system for studying the genetics of rapid adaptation, since it promises at the time to yield insights into the genetic pathways underlying the ecotype characteristics, including circadian and circalunar timing.

## Results

### *Clunio* ecotypes

Starting from field work in Northern Europe (Fig. 1A), we established one laboratory strain of the *Arctic ecotype* from Tromsø (Norway, Tro-tAR; see methods for strain nomenclature) and three laboratory strains of the *Baltic ecotype*, from Bergen (Norway, Ber-2AR), Sehlendorf (Germany; Seh-2AR) and Ar (Sweden; Ar-2AR). We also established a strain of the *Atlantic ecotype* from Bergen (Ber-1SL, sympatric with Ber-2AR) and used two existing *Atlantic ecotype* laboratory strains from Helgoland (Germany; He-1SL) and Port-en-Bessin (France; Por-1SL). We confirmed the identity of the ecotypes in the laboratory by the absence of a lunar rhythm in the *Baltic* and *Arctic ecotypes* (Fig. 1B-H), their circadian rhythm (Supplementary Fig. 1B-H) and their oviposition behavior (for details see Supplementary Note 1). The *Baltic ecotype* from Bergen (Ber-2AR, Fig. 1F) was found weakly lunar-rhythmic.

### Evolutionary history and species status

We sequenced the full nuclear and mitochondrial genomes of 168 field-caught individuals from six geographic sites, including the two sympatric population ecotypes from Bergen (Figure 1). Based on a set of 792,032 single nucleotide polymorphisms (SNPs), we first investigated population structure and evolutionary history by performing a principal component analysis (PCA; Fig. 2A-B) and testing for genetic admixture (Fig. 2C). We also constructed a haplotype network of complete mitochondrial genomes (Fig. 2D). There are several major observations that can be derived from these data.

**Figure 2.**
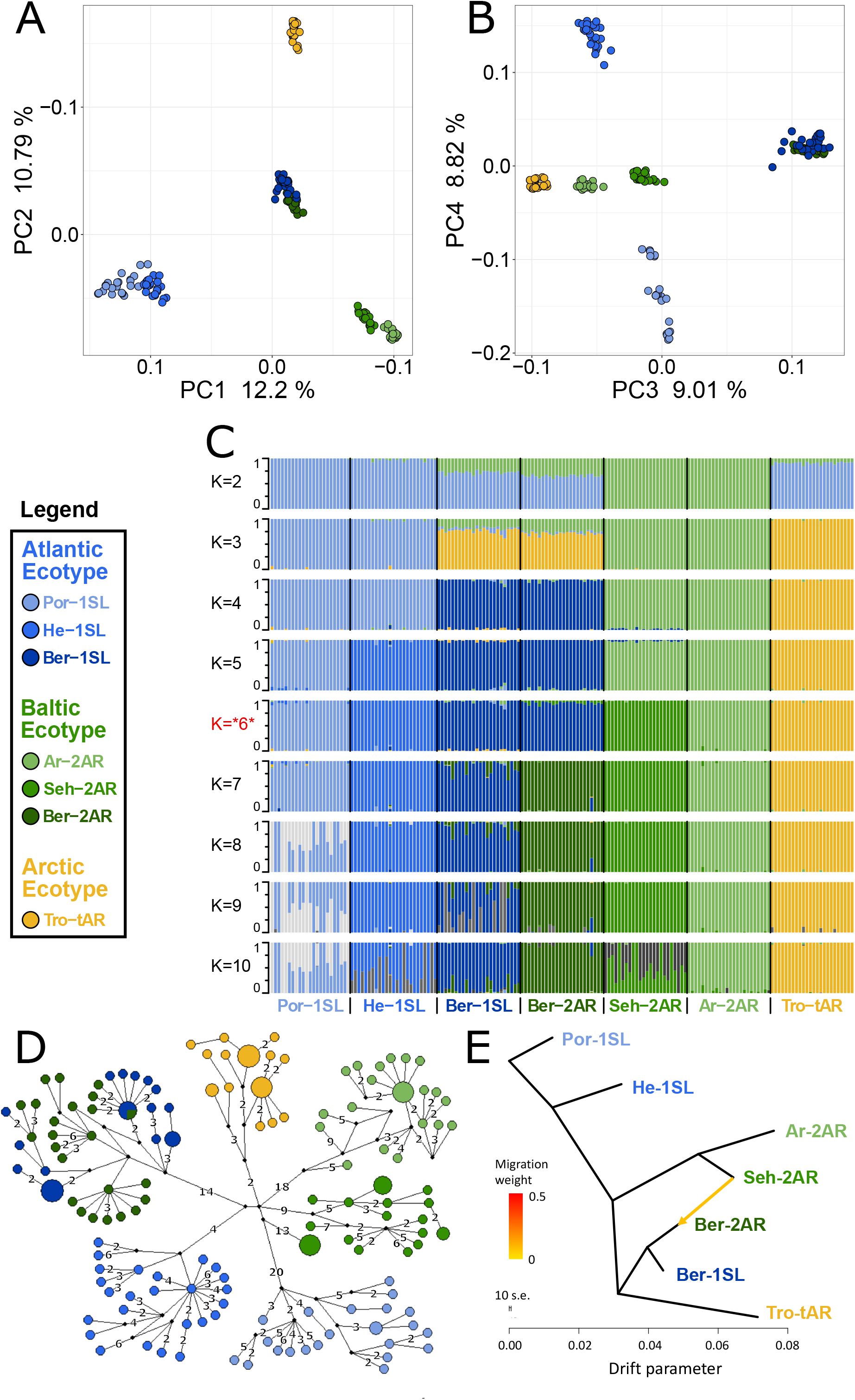
Genetic structure and evolutionary history of Northern European *Clunio* ecotypes. The analysis is based on 24 individuals from each population (23 for Por-1SL, 25 for He-1SL). **(A,B)** Principal Component Analysis (PCA) based on 792,032 SNPs separates populations by geographic location rather than ecotype. **(C)** ADMIXTURE analysis supports strong differentiation by geographic site (best K=6), but a notable genetic component from the Baltic Sea in the Bergen populations (see K=2 and 3). The Bergen populations are only separated at K=7 and then show a number of admixed individuals. **(D)** Haplotype network of full mitochondrial genomes reveals highly divergent clusters according to geographic site, but haplotype sharing between Ber-1SL and Ber-2AR. **(E)** Correlated allele frequencies indicate introgression from Seh-2AR into Ber-2AR.

First, there is strong geographic isolation between populations from different sites. In PCA, clusters are formed according to geography (Fig. 2A-B, Supplementary Fig. 2). Mitochondrial haplotypes are not shared and are diagnostically different between geographic sites (Fig. 2D). In ADMIXTURE, the optimal number of genetic groups is six (Fig. 2C, Supplementary Fig. 3), corresponding to the number of geographic sites, and there is basically no mixing between the six clusters (Fig. 2C; K=6). Finally, there is isolation by distance (IBD; Supplementary Figure 4).

Second, and much in contrast to the above, the sympatric ecotypes in Bergen are genetically very similar. In PCA they are not separated in the first four principal components (Fig. 2A-B) and they are the only populations that share mitochondrial haplotypes (Fig. 2D). In the ADMIXTURE analysis, they are only distinguished at K=7, a value larger than the optimal K. As soon as the two populations are distinguished, some individuals show signals of admixed origin (Fig. 2C; K=7), indicative of ongoing gene flow and incomplete reproductive isolation. These observations question the species status of *C. balticus*, which was based on the assumption of temporal isolation between these two populations^22^.

Third, the data suggest that after the ice age *Clunio* colonized northern Europe from a single source and expanded along two fronts into the Baltic Sea and into the high Arctic. The mitochondrial haplotype network expands from a single center, which implies a quick radiation from a colonizing haplotype (Fig. 2D; note that all sampled populations occur in areas under a former ice shield cover). There is also a large degree of shared nuclear polymorphisms. 34% of polymorphic SNPs are polymorphic in all seven populations and 93% are polymorphic in at least two populations (Supplementary Fig. 5). Separation of the Baltic Sea populations along PC1 and the Arctic population along PC2 (Fig. 2A), suggests that *Clunio* expanded into the high Arctic and into the Baltic Sea independently. Congruently, nucleotide diversity significantly decreases towards both expansion fronts (Supplementary Fig. 6, Supplementary Tab. 1). Postglacial establishment from a common source indicates that the *Baltic* and *Arctic ecotypes* have evolved their new local adaptations during this expansion time, i.e., in less than 10,000 years.

Fourth, ADMIXTURE analysis reveals that sympatric co-existence of the *Atlantic* and *Baltic ecotypes* in Bergen likely results from introgression of *Baltic ecotype* individuals into an existing *Atlantic ecotype* population. At K=2 and K=3 the two Baltic Sea populations Seh-2AR and Ar-2AR are separated from all other populations and the two Bergen populations Ber-2AR and Ber-1SL show a marked genetic contribution coming from these Baltic Sea populations (Fig. 2C). The Baltic genetic component is slightly larger for the *Baltic ecotype* Ber-2AR population than for the *Atlantic ecotype* Ber-1SL population. A statistical analysis via TreeMix detects predominantly an introgression from Seh-2AR into Ber-2AR (Fig. 2E, Supplementary Fig. 7), suggesting that genetic haplotypes that convey the *Baltic ecotype* have introgressed into a population of *Atlantic ecotype*, resulting in the now observed sympatric co-existence of *Baltic* and *Atlantic ecotypes* in Bergen.

While this observation adds to the evidence for a role of introgression in adaptation and early speciation processes ^25,26^, it also constitutes an excellent starting point for mapping loci that could be involved in the respective ecotype adaptations. Under the assumption that fast adaptation and ecotype formation are mostly based on frequency changes in standing variation, one expects that the respective haplotypes should be shared between ecotypes, especially when transferred via introgression. The following analysis is focussed on the *Atlantic* and *Baltic ecotypes*, represented by three populations each.

### Nuclear haplotype sharing

First, we reconstructed the genealogical relationship between 36 individuals (six from each population) in 50 kb windows (n=1,607) along the genome, followed by topology weighting. There are 105 possible unrooted tree topologies for six populations, and 46,656 possibilities to pick one individual from each population out of the set of 36. For each window along the genome, we assessed the relative support of each of the 105 population tree topologies by all 46,656 combinations of six individuals. We found that tree topologies change rapidly along the chromosomes (Fig. 3A; Supplementary Fig. 8; Supplementary Data 1). The tree topology obtained for the entire genome (Supplementary Fig. 9) dominates only in few genomic windows (Fig. 3A, black bars “Orig.”), while usually one or several other topologies account for more than 75% of the tree topologies (Fig. 3A, grey bars “Misc.”).

**Figure 3.**
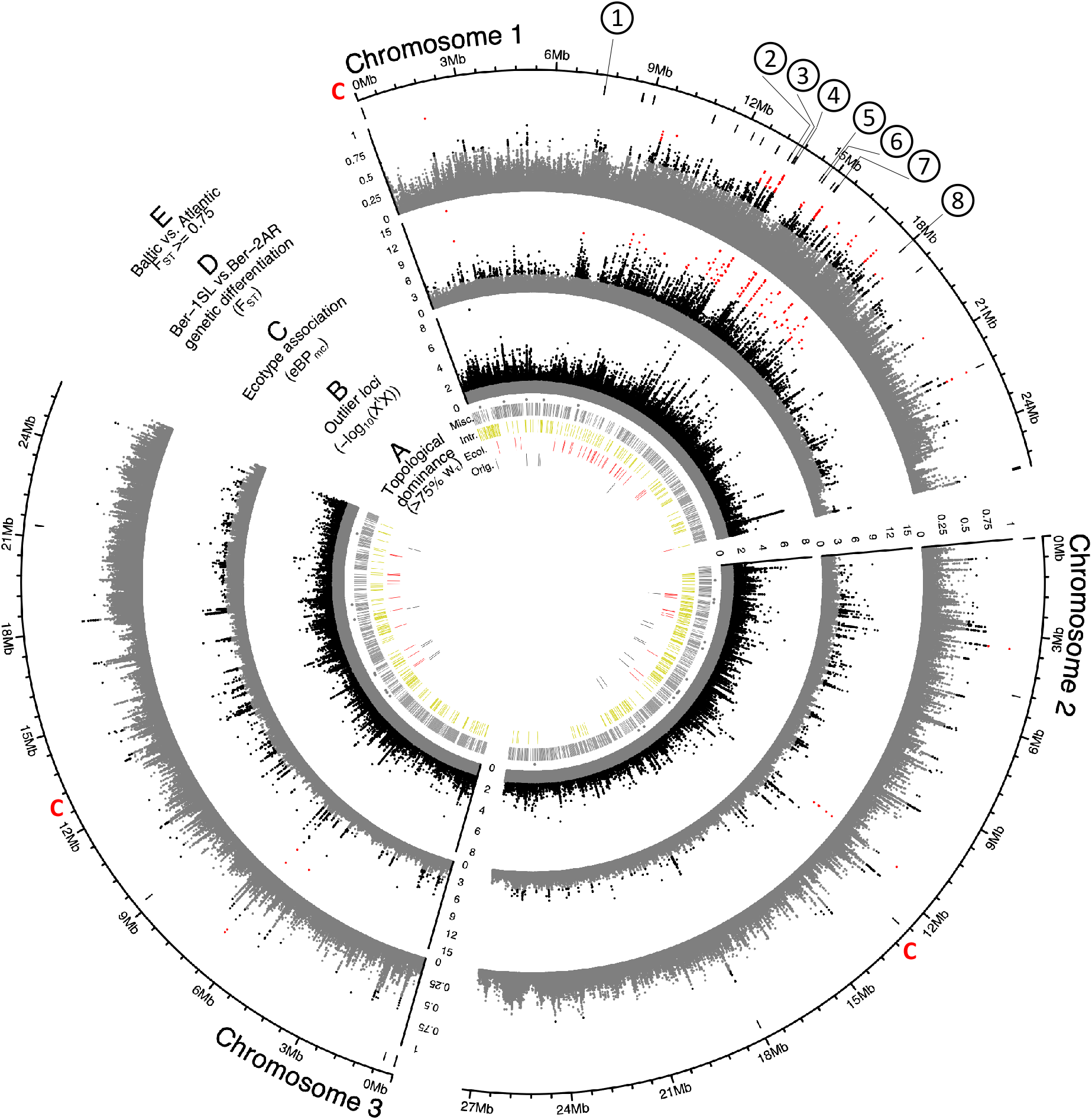
Genome screens for haplotype sharing and genotype-ecotype associations. (**A**) Topology weighting of phylogenetic trees for 36 individuals from the *Baltic* and *Atlantic ecotypes*, as obtained from 50 kb non-overlapping genomic windows. Windows were marked by a bar if they were dominated by one kind of topology (w_τ_ > 75%). Most windows are not dominated by the consensus population topology (“Orig.”; Suppl. Fig. 9), but by combinations of other topologies (“Misc.”). Windows dominated by topologies that separate the *Baltic* and *Atlantic ecotypes* (“Ecol.”) are mostly on chromosome 1. Windows consistent with introgression are found all over the genome (“Intr.”). (**B**) Distribution of outlier variants (SNPs and Indels) between the six *Baltic* and *Atlantic ecotype* populations, after global correction for population structure (X^t^X statistic). Values below the significance threshold (as obtained by subsampling) are plotted in grey. (**C**) Association of variant frequencies with *Baltic* vs. *Atlantic ecotype* (eBP_mc_). Values below the threshold of 3 (corresponding to p = 10^-3^) are given in grey, values above 10 are given in red. (**D**) Genetic differentiation (F_ST_) between the sympatric ecotypes in Bergen. Values above 0.5 are given in black, values above 0.75 in red. (**E**) The distribution of SNPs with F_ST_ >= 0.75 in the *Baltic* vs. *Atlantic ecotypes*. Circled numbers mark the location of the eight most differentiated loci (see Fig. 6). Centromeres of the chromosomes are marked by a red “C”.

Hardly ever do all combinations of six individuals follow a single population tree topology (Fig. 3A, stars), which implies that in most genomic windows some individuals do not group with their population. Taken together, this indicates a massive sharing of haplotypes across populations and high levels of incomplete lineage sorting, typical for such radiation situations. In such a highly mixed genomic landscape, it is close to impossible to separate signals of introgression from incomplete lineage sorting. Still, we highlighted genomic windows that are consistent with the detected introgression from the *Baltic ecotype* into both Bergen populations (Fig. 3A, yellow bars “Intr.”; all topologies grouping Por-1SL and He-1SL vs Ber-1SL, Ber-2AR, Seh-2AR and Ar-2AR). Regions consistent with introgression are scattered over the entire genome.

### Crosses and QTL mapping suggest a polygenic basis to lunar rhythmicity

As a first attempt to get hold of ecologically relevant loci, we assessed the genetic basis to lunar rhythmicity by crossing the lunar-arrhythmic Ber-2AR strain with the lunar-rhythmic Ber-1SL strain. Interestingly, the degree of lunar rhythmicity segregates within and between crossing families (Supplementary Fig. 10), suggesting a heterogeneous polygenic basis of lunar arrhythmicity. Genetic segregation in the cross implies that the weak rhythm observed in Ber-2AR is due to genetic polymorphism. The Ber-2AR strain seems to carry some lunar-rhythmic alleles, likely due to gene flow from the sympatric *Atlantic ecotype* (see Fig. 1C and results below).

For QTL mapping, we then picked a set of F2 families, which all go back to a single parental pair (BsxBa-F2 34; n=272 Supplementary Fig. 10). As a phenotype, we scored the degree of lunar rhythmicity for each individual as the number of days between the individual’s emergence and day 9 of the artificial turbulence cycle, which is the peak emergence day of the lunar-rhythmic strain (Supplementary Fig. 11). However, QTL analysis with various algorithms did not reveal any significant QTL for lunar rhythmicity (Fig. 4). As a control that in principle we had sufficient power for QTL mapping in this crossing family, we also mapped the sex determining locus, which was clearly detectable (Fig. 4). We can conclude that lunar rhythmicity must have a polygenic basis with many loci of small effects.

**Figure 4.**
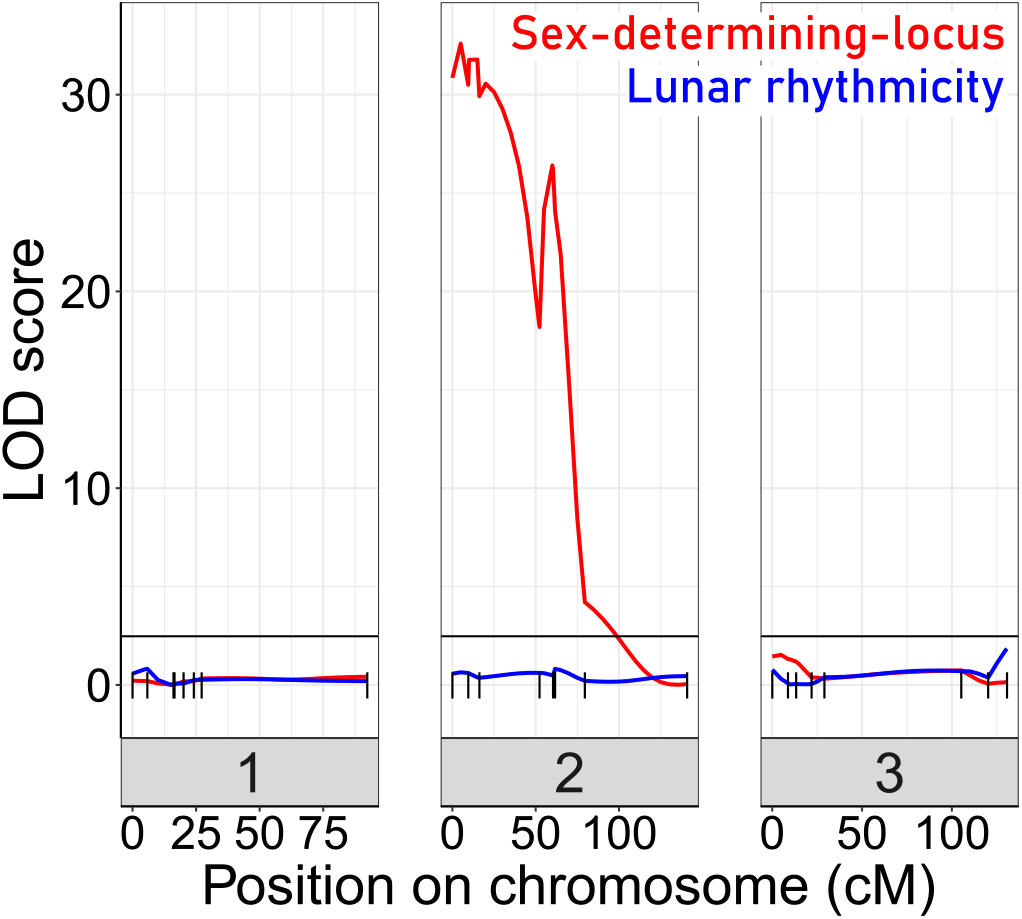
Quantitative Trait Locus (QTL) mapping for lunar rhythmicity and the sex determining locus. No significant QTLs can be detected for lunar rhythmicity, suggesting it is controlled by many loci of small effects. The sex-determining locus is clearly detectable, indicating that the is no general lack of power in the mapping panel.

### Genomic regions associated with ecotype formation

Given that QTL mapping for lunar rhythmicity suggested a polygenic basis to ecotype formation, we next applied three approaches to identify genomic regions associated with divergence between *Atlantic* and *Baltic ecotypes*.

First, genomic windows which are dominated by tree topologies that group populations according to ecotype were highlighted (Fig. 3A, red bars “Ecol.”).

Second, we screened all genetic variants (SNPs and indels; n=948,128) for those that are overly differentiated between the six populations after correcting for the neutral covariance structure across population allele frequencies (see Ω matrix, Supplementary Fig. 12A-B). Such variants can indicate local adaptation. At the same time, we tested for association of these variants with ecotype, as implemented in BayPass^27^. Overly differentiated variants (X^t^X statistic; Fig. 3B) and ecotype-associated variants ecotype (eBP_mc_; Fig. 3C) were detected all over the genome, but many were concentrated in the middle of the telocentric chromosome 1. Tests for association of variants with environmental variables such as sea surface temperature or salinity find fewer associated SNPs and no concentration on chromosome 1 (Supplementary Fig. 12D-E), confirming that the detected signals are not due to general genome properties, but are specific to the ecotypes.

Third, we expected that gene flow between the sympatric Ber-1SL and Ber-2AR populations would largely homogenize their genomes except for regions involved in ecological adaptation, which would be highlighted as peaks of genetic differentiation. The distributions of F_ST_ values in all pairwise population comparisons confirmed that genetic differentiation was particularly low in the Ber-1SL vs. Ber-2AR comparison (Supplementary Fig. 13 and 14). Pairwise differentiation between Ber-1SL and Ber-2AR (Fig. 3D) shows marked peaks on chromosome 1, most of which coincide with peaks in X^t^X and eBP_mc_. Notably, when assessing genetic differentiation of *Baltic* vs *Atlantic ecotype* (72 vs 72 individuals; Fig. 3E; Supplementary Fig. 15), there is not a single diagnostic variant (F_ST_ = 1), and even variants with F_ST_ ≥ 0.75 are very rare (n=63; Fig. 3E).

Genetic divergence (dxy), nucleotide diversity (*π*) and local linkage disequilibrium (*r^2^*) of the two Bergen populations do not show marked differences along or between chromosomes (Supplementary Fig. 16). The cluster of ecotype-associated variants on chromosome 1 overlaps with three large blocks of long-range linkage disequilibrium (LD; Supplementary Fig. 17). However, the boundaries of the LD blocks do not correspond to the ecotype-associated region and differ between populations. LD blocks are not ecotype-specific. Local PCA of the strongly ecotype associated region does not reveal patterns consistent with a chromosomal inversion or another segregating structural variant (Supplementary Fig. 18). Thus, there is no obvious link between the clustering of ecotype-associated loci and structural variation. Notably, genetic differentiation is not generally elevated in the ecotype-associated cluster on chromosome 1, as would be expected for a segregating structural variant, but drops to baseline levels in between ecotype-associated loci (Fig. 3D).

Taken together, these observations imply that numerous genomic loci – inside and outside the cluster on chromosome 1 – are associated with ecological adaptation and none of these are differentially fixed between ecotypes. In line with QTL mapping, this virtually excludes the possibility that only one or few loci have driven the new ecotype formation, suggesting instead that ecotype formation is based on a complex polygenic architecture.

### Support for adaptation from standing genetic variation

If, as assumed above, adaptation in these populations was indeed based mostly on standing genetic variation, one should find evidence in this in the data structure. We selected highly ecotype-associated SNPs (X^t^X > 1.152094, threshold obtained from randomized subsampling; eBP_mc_ > 3; n=3,976; Supplementary Fig. 19A) and assessed to which degree these alleles are shared between the studied populations and other populations across Europe. Allele sharing between the Bergen populations is likely due to ongoing gene flow, and hence Bergen populations were excluded from the analysis. In turn, allele sharing between the geographically isolated Seh-2AR, Ar-2AR, Por-1SL and He-1SL populations likely represents shared ancient polymorphism. Based on this comparison, we found that 82% of the ecotype-associated SNPs are polymorphic in both *Atlantic* and *Baltic ecotypes*, suggesting that the largest part of ecotype-associated alleles originates from standing genetic variation. We then retrieved the same genomic positions from published population resequencing data for *Atlantic ecotype* populations from Vigo (Spain) and St. Jean-de-Luz (Jean, southern France)^14^, an area that is potentially the source of postglacial colonization of all locations in this study. We found that 90% of the alleles associated with the Northern European ecotypes are also segregating in at least one of these southern populations, supporting the notion that adaptation in the North involves a re-assortment of existing standing genetic variation.

### Ecotypes differ mainly in the circadian clock and nervous system development

In the light of a polygenic architecture for ecotype formation, we can expect that usually several genes of a relevant physiological pathways are affected. This provides an ideal setting to identify the physiological processes underlying ecotype formation by enrichment analysis. Accordingly, we assessed how all ecotype-associated variants (SNPs and indels; X^t^X > 1.148764; eBP_mc_ > 3, n=4,741; Supplementary Fig. 19B) relate to *C. marinus’* genes. In a first step, we filtered the existing gene models in the CLUMA1.0 reference genome to those that are supported by transcript or protein evidence, have known homologues or PFAM domains, or were manually curated (filtered annotations provided in Supplementary Data 2; 15,193 gene models). Based on this confidence gene set, we then assessed the location of variants relative to genes, as well as the resulting mutational effects (SNPeff^28^; Supplementary Fig. 20; statistics in Supplementary Tab. 2). The vast majority of ecotype-specific variants are classified as intergenic modifier variants, suggesting that ecotype formation might primarily rely on regulatory mutations.

The ecotype-specific SNPs are found in and around 1,400 genes (Supplementary Data 3 and 4). We transferred GO terms from other species to the *Clunio* confidence annotations based on gene orthology (5,393 genes; see methods and Supplementary Data 5). GO term enrichment analysis suggests that ecological adaptation prominently involves the circadian clock, supported by three of the top four GO terms (Fig. 5A). In order to identify which genes drive GO term enrichment in the top 40 GO terms, we extracted the genes that harbour ecotype-associated SNPs (168 genes; Fig. 5B; Supplementary Table 3). We individually confirmed their gene annotations and associated GO terms. Clustering the resulting table by genes and GO terms reveals two dominant signatures (Fig. 5B). Many GO terms are associated with circadian timing and are driven by a small number of genes, which include almost all core circadian clock genes (Fig. 5B and C). As a second strong signal, almost half of the genes are annotated with biological processes involved in nervous system development (Fig. 5B and C). GO term enrichment is also found for ecdysteroid metabolism, imaginal disc development and gonad development (Fig. 5). These processes of pre-pupal development are expected to be under circalunar clock control. The fact that circalunar clocks are responsive to moonlight and water turbulence^9^ renders the finding of GO term enrichment for “auditory behaviour” and “phototaxis” interesting. Furthermore, many of the genes involved in nervous system development and sodium ion transport, also have GO terms that implicate them in light- and mechanoreceptor development, wiring or sensitivity (Supplementary Data 5). With the exception of “response to hypoxia” and possibly “sodium ion transmembrane transport”, there are very few GO terms that can be linked to the submerged larval habitat of the *Baltic ecotype*, which is usually low in salinity and can turn hypoxic in summer. There is a striking absence of GO terms involved in metabolic processes or immune response.

**Figure 5.**
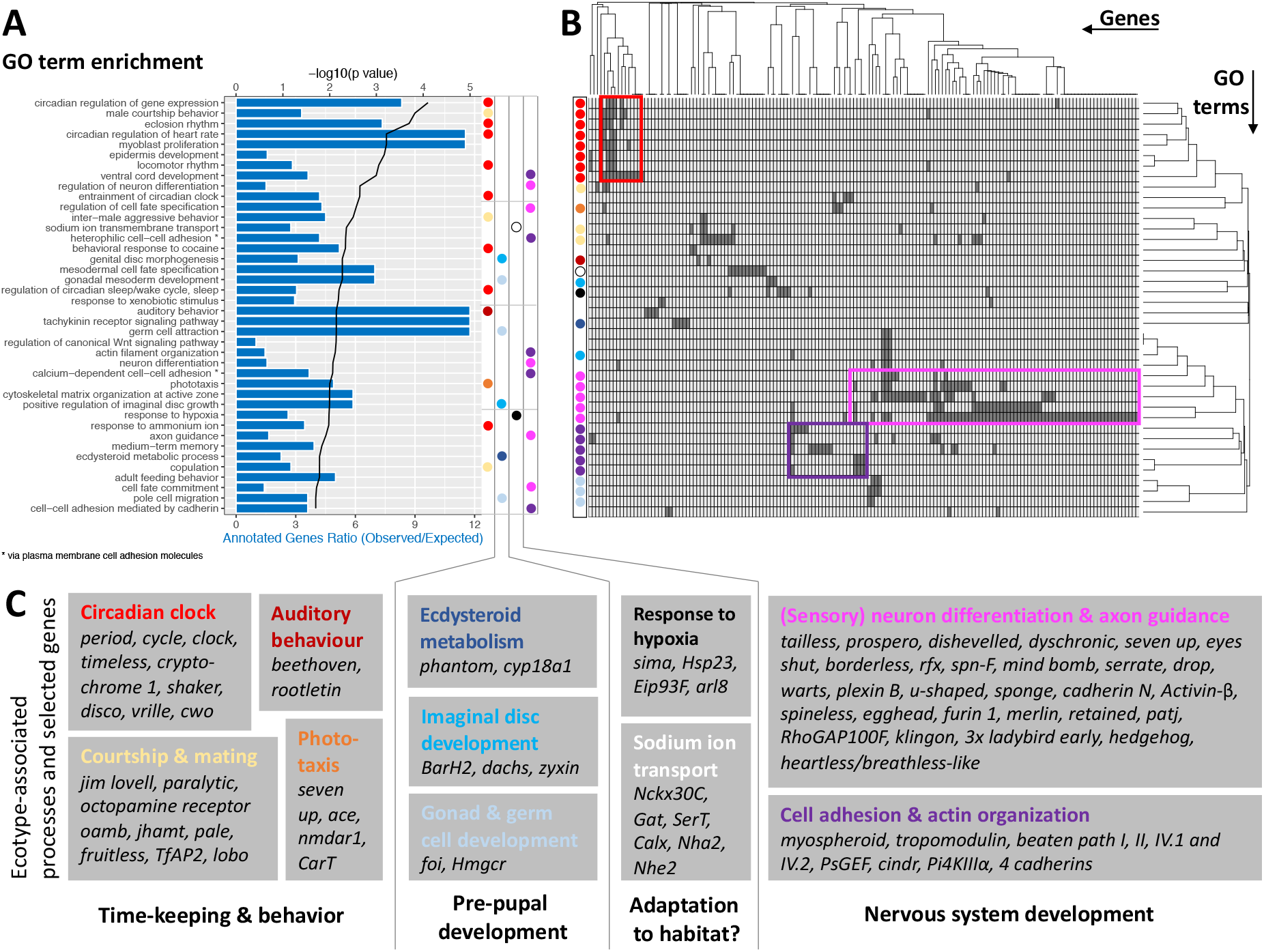
GO term analysis of ecotype associated SNPs. (**A**) The top 40 enriched GO terms are listed for the 1,400 genes that are found to be affected by ecotype-associated genetic variants (eBP_mc_ > 3). For each GO term the significance level (black line, top y-axis) and the observed-expected ratio of genes annotated to the respective GO term (blue bars, bottom y-axis) are given. (**B**) The top 40 GO terms are driven by 168 genes. Hierarchical clustering of genes and GO terms reveals major signals in the circadian clock and nervous system development (more details in Supplementary Table 3). (**C**) Most GO terms are consistent with the known ecotype differences and selected genes are highlighted for all of them. Notably, basically all core circadian clock genes are affected.

Taken together, the detected GO terms are highly consistent with the known ecotype differences and suggest that ecotypes are mainly defined by changes in the circadian clock and nervous system development. A previously unknown aspect of *Clunio* ecotype formation is highlighted by the GO terms “male courtship behaviour”, “inter-male aggression” and “copulation” (Fig. 5). These processes are subject to sexual selection and considered to evolve fast. They could in the long term entail assortative mating between ecotypes.

### Strongly differentiated loci correspond to GO-term enriched biological processes

While GO term analysis gives a broad picture of which processes have many genes affected by ecotype-associated SNPs, this does not necessarily imply that these genes and processes also show the strongest association with ecotype. Additionally, major genes might be missed because they were not assigned GO terms. As a second line of evidence, we therefore selected variants with the highest ecotype-association by increasing the eBP_mc_ cut-off to 10. This reduced the set of affected genes from 1,400 to 69 (Supplementary Data 6 and 7). Additionally, we only considered genes with variants that are strongly differentiated between the ecotypes (F_ST_ ≥ 0.75, compare Fig. 3E), leaving thirteen genes in eight distinct genomic regions (Fig. 6A; numbered in Fig. 3E). Two of these regions contain two genes each with no homology outside *C. marinus* (indicated by “NA”, Fig. 6A), confirming that GO term analysis missed major loci because of a lack of annotation. Three other regions contain the – likely non-visual – photoreceptor *ciliary Opsin 1* (*cOps1*)^29^, the transcription factor *longitudinals lacking* (*lola;* in fruit fly involved in axon guidance^30^ and photoreceptor determination^31^) and the nuclear receptor *tailless* (*tll*; in fruit fly involved in development of brain and eye^32^), underscoring that ecotype characteristics might involve differential light sensitivity. Interestingly, *tll* also affects development of the neuroendocrine centres involved in ecdysteroid production and adult emergence^33^. Even more, re-annotation of this genomic locus revealed that the neighbouring gene, which is also affected by ecotype specific variants, is the *short neuropeptide F receptor* (*sNPF-R*) gene. Among other functions, sNPF-R is involved in coupling adult emergence to the circadian clock^34^. Similarly, only 100 kb from *cOps1* there is the differentiated locus of *matrix metalloprotease 1 (Mmp1*), which is known to regulate circadian clock outputs via processing of the neuropeptide *pigment dispersing factor* (PDF)^35^. In both cases, the close genetic linkage could possibly form pre-adapted haplotypes and entail a concerted alteration of sensory and circadian functions in the formation of ecotypes. In the remaining two loci, *sox100B* is known to affect male gonad development^36^ and the *ecdysone-induced protein 93F* is involved in response to hypoxia in flies^37^, but was recently found to also affect reproductive cycles in mosquitoes^38^. In summary, only two out of the top 13 ecotype-associated genes were comprised in the top 40 GO terms (Fig. 6A). Nevertheless, all major biological processes detected in GO term analysis (Fig. 5) are also reflected in the strongly ecotype-associated loci (Fig. 6), giving a robust signal that circadian timing, sensory perception and nervous system development are underlying ecotype formation in *C. marinus*.

**Figure 6.**
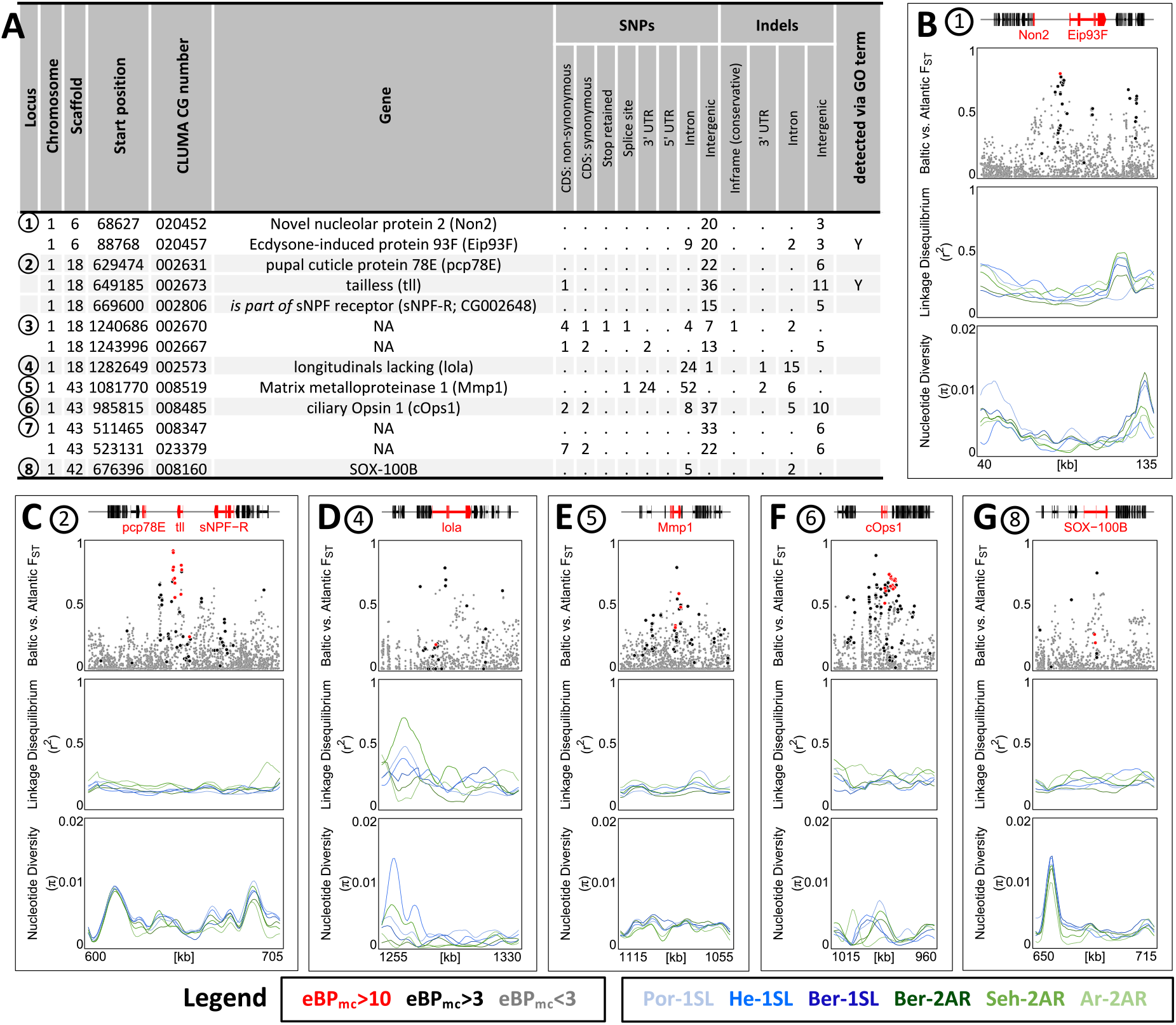
The 13 most differentiated ecotype-associated genes. (**A**) Loci with highly ecotype-associated variants were selected based on eBP_mc_ > 3 and F_ST_;(Baltic-Atlantic) > 0.75. There are 13 genes in eight distinct genomic loci. (**B-G**) An overview is given for the six loci with identified genes. In each panel, from top to bottom the sub-panels show the gene models, F_ST_ values of genetic variants in the region, local linkage disequilibrium (LD) and genetic diversity (*π*). F_ST_ values are coloured by ecotype association of the variant (red: eBP_mc_ > 10; black: 10 > eBP_mc_ > 3; grey: eBP_mc_ < 3). LD and genetic diversity are shown for the six populations independently, coloured-coded as in Fig. 1 and 2. The are no strong signatures of selection.

Finally, we assessed the top 13 strongly ecotype-associated loci for signatures of selective sweeps in genetic diversity and LD (Fig. 6B-G). Despite these loci being the most differentiated between ecotypes in the entire genome, there is at best a mild reduction in genetic diversity and a mild increase in LD (Fig. 6B-G). If selection acted on these loci, it must have been very soft, underscoring a history of polygenic adaptation from standing genetic variation and continued recombination.

## Discussion

The deep genomic analysis of *Clunio* populations has allowed to reconstruct a detailed biogeographic history of their different ecotypes. Most surprisingly, the two sympatric ecotypes that were considered different species turned out to be the most closely related populations in the sample, testifying the high power of ecology in generating differential adaptation in the face of gene flow. The data allowed further to infer a polygenic architecture for ecotype formation. We found that while ecotype-associated alleles differ in allele frequency, they are largely shared between ecotypes, which suggests that adaptation primarily involves standing genetic variation from many different loci. A similar re-use of existing regulatory variation has been found in ecotype formation in sticklebacks^39–41^ or mimicry in *Heliconius* butterflies^42^. However, while in *Heliconius* alleles are shared over large evolutionary distances via introgression, *Clunio* ecotypes diverged recently from a common source, as is illustrated by massive and genome-wide shared polymorphism. Combined with the observation that many genes from the same biological processes have ecotype-associated alleles, this draws a picture of polygenic adaptation, involving many pre-existing alleles with probably small phenotypic effects. Particularly for adaptation in circadian timing this scenario is highly plausible. The ancestral *Atlantic ecotype* comprises many genetically determined circadian timing types that are adapted to the local tides^11,12,14,43^. Existing genetic variants conveying emergence at dusk were likely selected or re-assorted to form the *Baltic ecotype’s* highly concentrated emergence at dusk.

Besides circadian timing, the ecotypes differ in circalunar timing and oviposition behavior. In our study the vast majority of GO terms and candidate genes is consistent with these functions, leaving little risk for evolutionary “story-telling” based on individual genes or GO terms^44^. We propose that good congruence between known phenotypic differences and detected biological processes could be a hallmark of polygenic adaptation, as only polygenic adaptation is expected to leave a footprint in many genes of the same ecologically relevant biological process.

Based on our genomic comparison of lunar-rhythmic and lunar-arrhythmic ecotypes, we propose three not mutually exclusive hypotheses on molecular pathways involved in the unknown circalunar clock. Firstly, *Clunio’s* circalunar clock is known to tightly regulate ecdysteroid-dependent development and maturation just prior to pupation^45^. Congruently, our screen identified ecotype-associated genes in the development of imaginal discs and genital discs, and in ecdysteroid metabolism. Lunar arrhythmicity may rely on an escape of these processes from circalunar clock control. Secondly, it has been hypothesized that circalunar clocks involve a circadian clock^46^ and such a mechanism has been experimentally confirmed in the midge *Pontomyia oceana^47^*. Thus, the overwhelming circadian signal in our data might be responsible for both circadian timing adaptations and the loss of circalunar rhythms. Thirdly, *Clunio’s* circalunar clock is synchronized with the external lunar cycle via moonlight, as well as tidal cycles of water turbulence and temperature^9^. Our data suggests that sensory receptor development, wiring or sensitivity might differ between ecotypes. Interestingly, some *Atlantic ecotype* populations are insensitive to specific lunar time cues, either moonlight or mechanical stimulation by the tides^11^. These pre-existing insensitivities may have been combined to form completely insensitive and hence lunar-arrhythmic ecotypes. This scenario would fit the general pattern of polygenic adaptation through a re-assortment of standing genetic variation, which emerges from our study.

In several species, genes involved in complex behavioral or ecological syndromes were found to be locked into supergenes by chromosomal inversions, e.g. in *Heliconius* butterfly mimicry^48^ or reproductive morphs of the ruff^49^. While we observe a clustering of ecotype-associated alleles in *Clunio*, there is no obvious connection to an underlying structural variant (SV). Possibly, the SV is so complex that it did not leave an interpretable genomic signal. Alternatively, *Clunio’s* long history of genome rearrangements^14^ may have resulted in a clustering of ecologically relevant loci without locking them into a single SV. Clustering could be stabilized by low recombination, consistent with the observed three LD blocks, which – while not ecotype-specific – all overlap with the differentiated region. Epistatic interactions between the clustered loci and co-adaptation of alleles might further reduce the fitness of recombinants and lead to a concerted response to selection. Such an interconnected adaptive cluster might allow for more flexible evolutionary responses than a single, completely linked supergene. Further studies will have to show whether such a genome architecture exists, whether it facilitates adaptation and whether it might itself be selected for.

## Methods

### Nomenclature of ecotypes

We expanded the existing naming convention of *C. marinus* timing types^43^ to also include *Baltic* and *Arctic ecotypes*. Names of populations and corresponding laboratory strains consist of an abbreviation for geographic origin followed by a code for the daily and lunar timing phenotypes. Daily phenotypes in this study are emergence during the first 12 hours after sunrise (“1”) or, emergence during the second 12 hours after sunrise (“2”) or emergence during every low tide (“t” for tidal rhythm). Lunar phenotypes in this study are either emergence during full moon and new moon low tides (“SL” for semi-lunar) or arrhythmic emergence (“AR”). As a consequence, the *Arctic ecotype* is “tAR”, the *Baltic ecotype* is “2AR” and the *Atlantic ecotype* populations in this study are all of timing type “1SL” (while other timing types exist within the *Atlantic ecotype^43^*).

### Fieldwork and sample collection

Field samples for genetic analysis and establishment of laboratory strains were collected in Sehlendorf (Seh, Germany), Ar (Sweden), Tromsø (Tro, Norway) and Bergen (Ber, Norway) during eight field trips in 2017 and 2018 (Supplementary Tab. 4). Field caught adult males for DNA extraction were directly collected in 99.98 % ethanol and stored at −20°C. Females are immobile and basically invisible in the field, unless found in copulation. Laboratory strains were established by catching copulating pairs in the field and transferring multiple fertilized egg clutches to the laboratory (Supplementary Tab. 4). Samples and laboratory strains of the sympatric ecotypes in Bergen were collected at the same location but at different daytime. Additional samples and laboratory strains from Helgoland (He, Germany) and Port-en-Bessin (Por, France) were collected and described earlier^14,43,50^, but had previously not been subject to whole genome sequencing of individuals.

### Laboratory culture and phenotyping of ecotypes

Laboratory strains were reared under standard conditions^51^ at 20°C with 16 h of light and 8 h of darkness. *Atlantic* and *Arctic ecotype* strains were kept in natural seawater diluted 1:1 with deionized water and fed with diatoms (*Phaeodactylum tricornutum*) and powdered nettles (*Urtica sp.*). The *Baltic ecotype* was kept in natural Sea water diluted 1:2 and fed with diatoms and powdered red algae (90%, *Delesseria spp*., 10% *Ceramium spp*., obtained from F. Weinberger and N. Stärck, GEOMAR, Kiel). For entrainment of the lunar rhythm all strains were provided with 12.4 h tidal cycles of water turbulence (mechanically induced vibrations produced by an unbalanced motor, 50 Hz, roughly 30 dB above background noise, 6.2 h on, 6.2 h off)^52,53^.

Assignment of strains to ecotypes was confirmed based on their phenotypes as recorded in laboratory culture. Oviposition behavior was assessed during standard culture maintenance: *Baltic ecotype* eggs are generally found submerged at the bottom of the culture vessel, *Atlantic* and *Arctic ecotype* eggs are always found floating on the water surface or on the walls of the culture vessel (see Supplementary Note 1). Daily emergence times were recorded in 1h intervals by direct observation (Seh-2AR, Ar-2AR) or with the help of a fraction collector^54^ (Ber-1SL, Ber-2AR, Tro-tAR, Por-1SL, He-1SL; Supplementary Fig. 1). Lunar emergence times were recorded by counting the number of emerged midges in the laboratory cultures every day over several months and summing them up over several tidal turbulence cycles. Emergence data for He-1SL was taken from^55^, emergence data for Por-1SL was taken from a manuscript in preparation (D Briševac, C Prakash, TS Kaiser).

### Crosses and Quantitative Trait Locus (QTL) mapping

The Ber-1SL and Ber-2AR laboratory strains were subject to single-pair crossing after their emergence rhythms were synchronized by keeping them in separate, time-shifted LD regimes. Several F1 families were raised independently and the F1 siblings were allowed to mate freely in order to obtain bulk F2 families. Emergence distributions were recorded by collecting all emerged adults every day.

Bulk crossing family 34 (n=272) was selected for QTL mapping. DNA of the two parents and the F2 progeny was extracted with the QuickExtract DNA Extraction Solution (Lucigen, QE0905T) according to manufacturer’s protocol with modifications. Instead of vortexing before incubation, a pestle was used to grind the sample. In order to identify scorable markers for the genetically very similar strains, the two parents of the cross were subject to whole-genome sequencing. Raw reads were processed through our genotyping pipeline (described below), resulting in 656,368 detected variants, 53,097 of which were diagnostic for the grandparents. Based on the list of diagnostic markers, we picked eight evenly spaced regions on each chromosome for amplicon sequencing (Supplementary Fig. 21, Supplementary Tab. 5). Multiplex PCR was performed with the QIAGEN Multiplex PCR Kit (206143) for each chromosome primer set separately (15 min at 95°C; 40 cycles of 94°C for 30 s, 57°C for 90 s and 72°C for 90 s; 72°C for 10 min). Sequencing libraries were prepared with standard Illumina protocols and sequenced with 150 bp paired-end on the Illumina HiSeq3000 sequencer by the Max Planck Genome Centre (Cologne, Germany). Sequencing reads from independent runs were first merged to one read file for forward and reverse reads. Those files were again subject to our genotyping pipeline (described below) with the exception that the mapped reads in SAM format undergone a coverage capping of 40 using a custom python script and the VCF was filtered for a minimum minor allele frequency of 0.25 (‘--maf 0.25’) and a maximum proportion of missing data per locus of 30% (‘--max-missing 0.7’). All markers that were not diagnostic for the parents were removed. In case genotypes differed within an amplicon, a consensus amplicon genotype was called by majority.

For QTL mapping the degree of rhythmicity of an individual was recorded as the number of days between its emergence day and day 9, which is the peak of emergence in the rhythmic strain (Ber-1SL). As this phenotype is not normally distributed (Kolmogorov-Smirnov test and the Shapiro-Wilk test of normality) it was treated as non-parametric. QTL scans were performed in the R package ‘qtl’ ^56^. The genome scans with different QTL models followed the calculation of conditional genotype probabilities in 5 cM distances and simulated genotypes from 64 imputations with the same distance. Both phenotypes (lunar rhythmicity, sex) were scanned for a single QTL model in the ‘scanone()’ function (method = “em”). Significance thresholds were set at top 5% of the logarithm of the odds (LOD) scores for 1000 permutations. The ‘stepwise.qtl()’ function for identification of multiple and interacting QTLs returned a null QTL model for lunar rhythmicity.

### DNA extraction and whole genome sequencing

For each of the seven populations, 24 field caught males (23 for Por-1SL, 25 for He-1SL) were subject to whole genome sequencing. DNA was extracted from entire individuals with a salting out method^57^ and amplified using the REPLI-g Mini Kit (QIAGEN) according to the manufacturer’s protocol with volume modifications (Supplementary Tab. 6). All samples were subject to whole genome shotgun sequencing at 15-20x target coverage on an Illumina HiSeq3000 sequencer with 150 bp paired-end reads. Library preparation and sequencing were performed by the Max Planck Genome Centre (Cologne, Germany) according to standard protocols. Raw sequence reads are deposited at ENA under Accession PRJEB43766.

### Sequence data processing, genotyping and SNP filtering

Raw sequence reads were trimmed for adapters and base quality using Trimmomatic v.0.38^58^ with parameters ‘ILLUMINACLIP:TruSeq3-PE-2.fa:2:30:10:8:true’, ‘LEADING:20’, ‘TRAILING:20’, ‘MINLEN:75’. Overlapping paired end reads were merged with PEAR v.0.9.10^59^, setting the minimum assembled sequence length to 75 bp and a capping quality score of 20. Assembled and unassembled reads were mapped with BWA-MEM^60^ to the nuclear reference genome^14^ (ENA accession GCA_900005825.1) and the mitochondrial reference genome (ENA accession CVRI01023763.1) of *C. marinus*. Mapped reads were sorted, indexed, filtered for mapping quality (-q 20) and transformed to BAM format with SAMtools v.1.9^61^. Read group information was added with the AddOrReplaceReadGroups.jar v.1.74 script from the Picard toolkit (http://picard.sourceforge.net/) ^62^.

For the nuclear genome, SNPs and insertion-deletion (indel) genotypes were called using GATK v.3.8-0-ge9d806836^63^. After initial genotype calling with the GATK HaplotypeCaller and the parameter ‘-stand_call_conf 30’, base qualities were recalibrated with the GATK BaseRecalibrator with ‘-knownSites’ and genotype calling was repeated on the recalibrated BAM files to obtain the final individual VCF files. Individual VCF files were combined using GATK GenotypeGVCFs. SNP and indel genotypes were filtered with VCFtools v.0.1.14 ^64^ to keep only biallelic polymorphisms (--max-alleles 2), with a minimum minor allele frequency of 0.02 (-- maf 0.02), a minimum genotype quality of 20 (--minQ 20) and a maximum proportion of missing data per locus of 40% (--max-missing 0.6), resulting in 792,032 SNPs and 156,096 indels over the entire set of 168 individuals. For certain analyses indels were excluded with VCFtools (‘--remove-indels’).

Reads mapped to the mitochondrial genome were transformed into mitochondrial haplotypes as described in ^65^.

### Population genomic analyses

Mitochondrial haplotype networks were calculated using the Median-Joining algorithm^66^ with Network v.10.1.0.0 (fluxus-engineering.com).

Nuclear SNP genotypes were converted to PLINK format with VCFtools. SNPs were LD pruned with PLINK v.1.90b4^67^ and parameters ‘--indep-pairwise 50 10 0.5’ as well as ‘--chr-set 3 no-xy no-mt --nonfounders’. Principal Component Analysis (PCA) was performed in PLINK using the option ‘--pca’ with the options default settings. The pruned BED file from PLINK was used as input to ADMIXTURE v.1.3.0^68^, with which we assessed a series of models for K=1 to K=10 genetic components, as well as the corresponding the cross-validation error (‘--cv’). Migration was further tested by converting the SNP data to TreeMix format with the *vcf2tree-mix.sh* script^69^ and running *TreeMix 1.13^70^* with default parameters and the southernmost Por-1SL as root population.

Population estimates along the chromosomes were calculated in 100 kb overlapping sliding-windows with 10 kb steps. Nucleotide diversity (*π*) was calculated for SNPs with VCFtools ‘--window-pi’. For the genome-wide average, calculations were repeated with 200kb non-overlapping windows. Linkage disequilibrium (LD; as r^2^) was calculated in VCFtools with ‘-- geno-r2’. Local LD was calculated with ‘--ld-window-bp 500’. Preliminary tests showed that local LD decays within a few hundred base pairs (Supplementary Fig. 22). For long range LD minor allele frequency was filtered to 0.2 (‘--maf 0.2’, resulting in 335,800 SNPs), only values larger 0.5 were allowed with ‘--min-r2 0.5’ and the ‘--ld-window-bp 500’ filter was removed. Pairwise F_ST_ was calculated with VCFtools ‘--weir-fst-pop’ option per SNP and in sliding windows. For calculation of genetic divergence (dxy), allele frequencies were extracted with VCFtools ‘--freq’ and dxy was estimated from allele frequencies according to ^71^.

### Phylogenomics and topology weighting

Nuclear genome phylogeny was calculated for a random set of six individuals from each population, without Tro-tAR (n=36). For windowed phylogenies, the VCF file was subset into non-overlapping 50 kb windows using VCFtools ‘-- from-bp --to-bp’. SNP genotypes were transformed into FASTA alignments of only informative sites with the vcf2phylip.py v.2.3 script^72^ and parameters ‘-m 1 -p -f’. Heterozygous genotypes were represented by the respective IUPAC code for both bases. Whole genome and windowed phylogenies were calculated with IQ-TREE v.1.6.12 ^73^ using the parameters ‘-st DNA -m MFP -keep-ident -redo’ for the windowed and ‘-st DNA -m MFP -keep-ident -bb 1000 -bnni -nt 10 - redo’ for the whole genome phylogenies. Topology weighting was performed on the windowed phylogenies with TWISST ^74^ and the parameter ‘--method complete’.

### Association analysis

Population-based association between genetic variants (SNPs and Indels) and ecotype, as well as environmental variables (Supplementary Tab. 7) was assessed in BayPass v.2.2 ^27^. Allele counts were obtained with VCFtools option ‘--counts’. Analyzed covariates were ecotype, sea surface salinity (obtained from ^75^) and average water temperature of the year 2020 (obtained from weather-atlas.com, accessed 27.04.2020; 16:38), as given in Supplementary Tab. 7. BayPass was run with the MCMC covariate model. BayPass corrects for population structure via Ω dissimilarity matrices, then calculates the X^t^X statistics and finally assesses the approximate Bayesian p value of association (eBP_mc_). To obtain a significance threshold for X^t^X values, the data was randomly subsampled (100,000 genetic variants) and re-analyzed with the standard covariate model, as implemented in baypass_utils.R. All analyses we performed in three replicates (starting seeds 5001, 24306 and 1855) and the median is shown.

### SNP effects and GO term enrichment analysis

Gene annotations to the CLUMA1.0 reference genome ^14^ were considered reliable if they fulfilled one of three criteria: 1) Identified ortholog in UniProtKB/Swiss-Prot or non-redundant protein sequences (nr) at NCBI or PFAM domain, as reported in ^14^. 2) Overlap of either at least 20% with mapped transcript data or 40% with mapped protein data, as reported in ^14^. 3) Manually annotated. This resulted in a 15,193 confidence genes models. The location and putative effects of the SNPs and indels relative to these confidence gene models were annotated using SnpEff 4.5 ^28^ (build 2020-04-15 22:26, non-default parameter ‘-ud 0’). Gene Ontology (GO) terms were annotated with emapper-2.0.1. ^76^ from the eggNOG 5.0 database^77^, using DIAMOND ^78^, BLASTP e-value <1e^-10^ and subject-query alignment coverage of >60%. Conservatively, we only transferred GO terms with “non-electronic” GO evidence from best-hit orthologs restricted to an automatically adjusted per-query taxonomic scope, resulting in 5,393 *C. marinus* gene models with GO term annotations. Enrichment of “Biological Process” GO terms in the genes associated with ecotype-specific polymorphisms was assessed with the weight01 Fisher’s exact test implemented in topGO ^79^ (version 2.42.0, R version 4.0.3).

### Figure preparation

Figures were prepared in R^80^. Data were handled with the ‘data.table’^81^ and ‘plyr’^82^ packages. The map of Europe was generated using the packages ‘ggplot2’^83^ and ‘ggrepel’ ^84^, ‘maps’^85^ and ‘mapdata’^86^. The map was taken from the CIA World DataBank II (http://www.evl.uic.edu/pape/data/WDB/). Circular plots were prepared using the R package ‘circlize’^87^. Multiple plots were combined in R using the package ‘Rmisc’^88^. The graphical editing of the whole genome phylogeny was done in Archeopteryx (http://www.phylosoft.org/ar-chaeopteryx)^89^. Final figure combination and graphical editing of the raw plot files was done in *Inkscape*. Neighbor Joining trees of the omega statistic distances from BayPass were created with the R package ‘ape’.^90^ In all plots the order and orientation of scaffolds within the chromosomes follows the published genetic linkage map^14^.

## Data availability

Sequencing data for field samples and mapping families are deposited at ENA under Accession PRJEB43766. Filtered genome annotations are given in Supplementary Dataset 2.

## Acknowledgements

For field work we obtained logistic support from the Ar Research Station (Uppsala University), the Marine Biological Station Espegrend (University of Bergen), Even Jørgensen (The Arctic University of Norway, Tromsø) and Florian Weinberger and Nadja Stärck (GEOMAR Helmholtz Centre for Ocean Research Kiel). We thank Jürgen Reunert, Kerstin Schäfer and Susanne Mentz for technical assistance, as well as all members of the MPRG “Biological Clocks” for discussion and support. Diethard Tautz and Julien Dutheil critically read the manuscript. Whole genome sequencing was performed at the Max Planck Genome Center (Cologne) with financial support from the Max Planck Society. This work was funded by the Max Planck Society through the Max Planck Research Group “Biological Clocks” and a sequencing grant. The work was further funded by the European Research Council (ERC) under the Horizon 2020 research and innovation program with an ERC Starting Grant (Grant agreement 802923) awarded to TSK.

## Author contributions

NF performed field work, laboratory work, QTL mapping, population genomic analyses and association analysis, prepared the figures and participated in drafting the manuscript. CP analyzed SNP effects and GO term enrichment and participated in figure preparation. TSK conceived, designed and supervised the study, participated in data analysis and figure preparation and wrote the manuscript. All authors approved of the final manuscript.

## Competing interests

There are no competing interests.

## Notes

### Competing Interest Statement

The authors have declared no competing interest.

### Summary of Updates

QTL mapping added (new Figure 4 and new section); text edited; supplemental files updated.

## References

1 Wu, C.-I. & Ting, C.-T. Genes and speciation. Nature Reviews Genetics 5, 114–122 (2004).

2 Phadnis, N. & Orr, H. A. A single gene causes both male sterility and segregation distortion in Drosophila hybrids. science 323, 376–379 (2009).

3 Nosil, P. & Schluter, D. The genes underlying the process of speciation. Trends in ecology & evolution 26, 160–167 (2011).

4 Hof, A. E. et al. The industrial melanism mutation in British peppered moths is a transposable element. Nature 534, 102–105 (2016).

5 Sella, G. & Barton, N. H. Thinking about the evolution of complex traits in the era of genome-wide association studies. Annual review of genomics and human genetics 20 (2019).

6 Boyle, E. A., Li, Y. I. & Pritchard, J. K. An expanded view of complex traits: from polygenic to omnigenic. Cell 169, 1177–1186 (2017).

7 Barton, N. H. The “New Synthesis”. Proceedings of the National Academy of Sciences 119, e2122147119, doi:doi:10.1073/pnas.2122147119 (2022).

8 Barghi, N., Hermisson, J. & Schlötterer, C. Polygenic adaptation: a unifying framework to understand positive selection. Nature Reviews Genetics 21, 769–781 (2020).

9 Neumann, D. in Annual, Lunar, and Tidal Clocks (eds Hideharu Numata & Barbara Helm) Ch. 1, 3–24 (Springer Japan, 2014).

10 Andreatta, G. & Tessmar-Raible, K. The still dark side of the moon: molecular mechanisms of lunar-controlled rhythms and clocks. J Mol Biol 432, 3525–3546 (2020).

11 Kaiser, T. S. in Annual, Lunar, and Tidal Clocks (eds Hideharu Numata & Barbara Helm) Ch. 7, 121–141 (Springer Japan, 2014).

12 Neumann, D. Genetic adaptation in emergence time of *Clunio* populations to different tidal conditions. Helgoländer wissenschaftliche Meeresuntersuchungen 15, 163–171 (1967).

13 Kaiser, T. S., Neumann, D. & Heckel, D. G. Timing the tides: Genetic control of diurnal and lunar emergence times is correlated in the marine midge *Clunio marinus*. BMC Genetics 12, 49, doi:10.1186/1471-2156-12-49 (2011).

14 Kaiser, T. S. et al. The genomic basis of circadian and circalunar timing adaptations in a midge. Nature 540, 69–73, doi:10.1038/nature20151 (2016).

15 Kaiser, T. S. & Heckel, D. G. Genetic Architecture of Local Adaptation in Lunar and Diurnal Emergence Times of the Marine Midge *Clunio marinus* (Chironomidae, Diptera). PLoS ONE 7, e32092, doi:10.1371/journal.pone.0032092 (2012).

16 Remmert, H. Ökologische Untersuchungen über die Dipteren der Nord-und Ostsee. Archiv für Hydrobiologie 51, 1–53 (1955).

17 Endraß, U. Physiologische Anpassungen eines marinen Insekts. I. Die zeitliche Steuerung der Entwicklung. Marine Biology 34, 361–368 (1976).

18 Palmén, E. & Lindeberg, B. The marine midge, Clunio marinus Hal. (Dipt., Chironomidae), found in brackish water in the northern Baltic. Internationale Revue der gesamten Hydrobiologie und Hydrogeographie 44, 383–393 (1959).

19 Neumann, D. & Honegger, H. W. Adaptations of the Intertidal Midge Clunio to Arctic Conditions. Oecologia 3, 1–13 (1969).

20 Pflüger, W. & Neumann, D. Die Steuerung einer gezeitenparallelen Schlüpfrhythmik nach dem Sanduhr-Prinzip. Oecologia 7, 262–266 (1971).

21 Endraß, U. Physiologische Anpassungen eines marinen Insekts. II. Die Eigenschaften von schwimmenden und absinkenden Eigelegen. Marine Biology 36, 47–60 (1976).

22 Heimbach, F. Sympatric species, Clunio marinus Hal. and Cl. balticus n. sp. (Dipt., Chironomidae), isolated by differences in diel emergence time. Oecologia 32, 195–202 (1978).

23 Patton, H. et al. Deglaciation of the Eurasian ice sheet complex. Quaternary Science Reviews 169, 148–172 (2017).

24 Hofmann, W. & Winn, K. The littorina transgression in the Western Baltic Sea as indicated by subfossil Chironomidae (Diptera) and Cladocera (Crustacea). International Review of Hydrobiology 85, 267–291 (2000).

25 Edelman, N. B. & Mallet, J. Prevalence and adaptive impact of introgression. Annual Review of Genetics 55, 265–283 (2021).

26 Baack, E. J. & Rieseberg, L. H. A genomic view of introgression and hybrid speciation. Current opinion in genetics & development 17, 513–518 (2007).

27 Gautier, M. Genome-wide scan for adaptive divergence and association with population-specific covariates. Genetics 201, 1555–1579 (2015).

28 Cingolani, P. et al. A program for annotating and predicting the effects of single nucleotide polymorphisms, SnpEff: SNPs in the genome of *Drosophila melanogaster* strain *w*^1118^; *iso*-2; *iso-3*. Fly 6, 80–92, doi:10.4161/fly.19695 (2012).

29 Velarde, R. A., Sauer, C. D., Walden, K. K. O., Fahrbach, S. E. & Robertson, H. M. Pteropsin: A vertebrate-like non-visual opsin expressed in the honey bee brain. Insect Biochemistry and Molecular Biology 35, 1367–1377 (2005).

30 Crowner, D., Madden, K., Goeke, S. & Giniger, E. Lola regulates midline crossing of CNS axons in Drosophila. Development 129, 1317–1325 (2002).

31 Zheng, L. & Carthew, R. W. Lola regulates cell fate by antagonizing Notch induction in the Drosophila eye. Mechanisms of Development 125, 18–29 (2008).

32 Suzuki, T. & Saigo, K. Transcriptional regulation of atonal required for Drosophila larval eye development by concerted action of eyes absent, sine oculis and hedgehog signaling independent of fused kinase and cubitus interruptus. Development 127, 1531–1540 (2000).

33 de Velasco, B. et al. Specification and development of the pars intercerebralis and pars lateralis, neuroendocrine command centers in the Drosophila brain. Developmental Biology 302, 309–323 (2007).

34 Selcho, M. et al. Central and peripheral clocks are coupled by a neuropeptide pathway in Drosophila. Nature Communications 8, 15563, doi:10.1038/ncomms15563 (2017).

35 Depetris-Chauvin, A. et al. Mmp1 Processing of the PDF Neuropeptide Regulates Circadian Structural Plasticity of Pacemaker Neurons. PLOS Genetics 10, e1004700, doi:10.1371/journal.pgen.1004700 (2014).

36 Nanda, S. et al. Sox100B, a Drosophila group E Sox-domain gene, is required for somatic testis differentiation. Sexual Development 3, 26–37 (2009).

37 Lee, S.-J., Feldman, R. & O’Farrell, P. H. An RNA interference screen identifies a novel regulator of target of rapamycin that mediates hypoxia suppression of translation in Drosophila S2 cells. Molecular Biology of the Cell 19, 4051–4061 (2008).

38 Wang, X. et al. The ecdysone-induced protein 93 is a key factor regulating gonadotrophic cycles in the adult female mosquito Aedes aegypti. Proceedings of the National Academy of Sciences 118(2021).

39 Chan, Y. F. et al. Adaptive evolution of pelvic reduction in sticklebacks by recurrent deletion of a Pitx1 enhancer. Science 327, 302–305 (2010).

40 Kingman, G. A. R. et al. Predicting future from past: The genomic basis of recurrent and rapid stickleback evolution. bioRxiv (2020).

41 Verta, J.-P. & Jones, F. C. Predominance of cis-regulatory changes in parallel expression divergence of sticklebacks. Elife 8, e43785 (2019).

42 Edelman, N. B. et al. Genomic architecture and introgression shape a butterfly radiation. Science 366, 594–599, doi:10.1126/science.aaw2090 (2019).

43 Kaiser, T. S., von Haeseler, A., Tessmar-Raible, K. & Heckel, D. G. Timing strains of the marine insect Clunio marinus diverged and persist with gene flow. Molecular Ecology 30, 1264–1280, doi:https://doi.org/10.1111/mec.15791(2021).

44 Pavlidis, P., Jensen, J. D., Stephan, W. & Stamatakis, A. A Critical Assessment of Storytelling: Gene Ontology Categories and the Importance of Validating Genomic Scans. Molecular Biology and Evolution 29, 3237–3248, doi:Doi 10.1093/Molbev/Mss136 (2012).

45 Neumann, D. & Spindler, K. D. Circasemilunar Control of Imaginal Disk Development in Clunio marinus - Temporal Switching Point, Temperature-Compensated Developmental Time and Ecdysteroid Profile. Journal of Insect Physiology 37, 101–109 (1991).

46 Bünning, E. & Müller, D. Wie messen Organismen lunare Zyklen? Zeitschrift für Naturforschung 16, 391–395 (1962).

47 Soong, K. & Chang, Y.-H. Counting Circadian Cycles to Determine the Period of a Circasemilunar Rhythm in a Marine Insect. Chronobiology International 29, 1329–1335, doi:10.3109/07420528.2012.728548 (2012).

48 Joron, M. et al. Chromosomal rearrangements maintain a polymorphic supergene controlling butterfly mimicry. Nature 477, 203–U102, doi:10.1038/nature10341 (2011).

49 Küpper, C. et al. A supergene determines highly divergent male reproductive morphs in the ruff. Nature Genetics 48, 79–83 (2016).

50 Kaiser, T. S., Neumann, D., Heckel, D. G. & Berendonk, T. U. Strong genetic differentiation and postglacial origin of populations in the marine midge *Clunio marinus* (Chironomidae, Diptera). Molecular Ecology 19, 2845–2857, doi:10.1111/j.1365-294X.2010.04706.x (2010).

51 Neumann, D. Die lunare und tägliche Schlüpfperiodik der Mücke *Clunio* - Steuerung und Abstimmung auf die Gezeitenperiodik. Zeitschrift für Vergleichende Physiologie 53, 1–61 (1966).

52 Neumann, D. & Heimbach, F. in Cyclic Phenomena in Marine Plants and Animals (eds E. Naylor & R.G. Hartnoll) 423–433 (Pergamon Press, 1979).

53 Neumann, D. Entrainment of a Semilunar Rhythm by Simulated Tidal Cycles of Mechanical Disturbance. Journal of Experimental Marine Biology and Ecology 35, 73–85 (1978).

54 Honegger, H. W. An automatic device for the investigation of the rhythmic emergence pattern of Clunio marinus. International Journal of Chronobiology 4, 217–221 (1977).

55 Neumann, D. Die zeitliche Programmierung von Tieren auf periodische Umweltbedingungen. Rheinisch-Westfälische Akademie der Wissenschaften, Natur-Ingenieur-und Wirtschaftswissenschaften, Vortraege, 31–62 (1983).

56 Broman, K. W., Wu, H., Sen, Ś. & Churchill, G. A. R/qtl: QTL mapping in experimental crosses. bioinformatics 19, 889–890 (2003).

57 Reineke, A., Karlovsky, P. & Zebitz, C. P. W. Preparation and purification of DNA from insects for AFLP analysis. Insect Molecular Biology 7, 95–99 (1998).

58 Bolger, A. M., Lohse, M. & Usadel, B. Trimmomatic: a flexible trimmer for Illumina sequence data. Bioinformatics 30, 2114–2120 (2014).

59 Zhang, J., Kobert, K., Flouri, T. & Stamatakis, A. PEAR: a fast and accurate Illumina Paired-End reAd mergeR. Bioinformatics 30, 614–620 (2014).

60 Li, H. Aligning sequence reads, clone sequences and assembly contigs with BWA-MEM. arXiv preprint arXiv:1303.3997 (2013).

61 Li, H. et al. The Sequence Alignment/Map format and SAMtools. Bioinformatics 25, 2078–2079, doi:10.1093/bioinformatics/btp352 (2009).

62 DePristo, M. A. et al. A framework for variation discovery and genotyping using next-generation DNA sequencing data. Nature genetics 43, 491 (2011).

63 McKenna, A. et al. The Genome Analysis Toolkit: A MapReduce framework for analyzing next-generation DNA sequencing data. Genome Research 20, 1297–1303, doi:10.1101/gr.107524.110 (2010).

64 Danecek, P. et al. The variant call format and VCFtools. Bioinformatics 27, 2156–2158 (2011).

65 Fuhrmann, N. & Kaiser, T. S. The importance of DNA barcode choice in biogeographic analyses—a case study on marine midges of the genus Clunio. Genome, 1–11 (2020).

66 Bandelt, H. J., Forster, P. & Rohl, A. Median-joining networks for inferring intraspecific phylogenies. Molecular Biology and Evolution 16, 37–48 (1999).

67 Chang, C. C. et al. Second-generation PLINK: rising to the challenge of larger and richer datasets. Gigascience 4, s13742-13015-10047-13748 (2015).

68 Alexander, D. H. & Lange, K. Enhancements to the ADMIXTURE algorithm for individual ancestry estimation. BMC Bioinformatics 12, 246, doi:10.1186/1471-2105-12-246 (2011).

69 Ravinet, M. <https://github.com/speciationgenomics/scripts/blob/master/vcf2treemix.sh> (last accessed 16th April 2021).

70 Pickrell, J. & Pritchard, J. Inference of population splits and mixtures from genome-wide allele frequency data. Nature Precedings, 1–1 (2012).

71 Delmore, K. E. et al. Genomic analysis of a migratory divide reveals candidate genes for migration and implicates selective sweeps in generating islands of differentiation. Molecular Ecology 24, 1873–1888 (2015).

72 Ortiz, E. vcf2phylip v2. 0: convert a VCF matrix into several matrix formats for phylogenetic analysis. URL https://doi.org/10.5281/zenodo 2540861 (2019).

73 Nguyen, L.-T., Schmidt, H. A., Von Haeseler, A. & Minh, B. Q. IQ-TREE: a fast and effective stochastic algorithm for estimating maximum-likelihood phylogenies. Molecular Biology and Evolution 32, 268–274 (2015).

74 Martin, S. H. & Van Belleghem, S. M. Exploring evolutionary relationships across the genome using topology weighting. Genetics 206, 429–438 (2017).

75 Hordoir, R. et al. Nemo-Nordic 1.0: a NEMO-based ocean model for the Baltic and North seas–research and operational applications. Geoscientific Model Development 12, 363–386 (2019).

76 Huerta-Cepas, J. et al. Fast genome-wide functional annotation through orthology assignment by eggNOG-mapper. Molecular Biology and Evolution 34, 2115–2122 (2017).

77 Huerta-Cepas, J. et al. eggNOG 5.0: a hierarchical, functionally and phylogenetically annotated orthology resource based on 5090 organisms and 2502 viruses. Nucleic Acids Research 47, D309–D314 (2019).

78 Buchfink, B., Xie, C. & Huson, D. H. Fast and sensitive protein alignment using DIAMOND. Nature Methods 12, 59–60 (2015).

79 Alexa, A. & Rahnenfuhrer, J. topGO: enrichment analysis for gene ontology. R package version 2, 2010 (2010).

80 Crawley, M. J. The R Book. (John Wiley & Sons Ltd., 2007).

81 Dowle, M. et al. Package ‘data.table’ version 1.13.2. Extension of ‘data. frame (2020).

82 Wickham, H. The split-apply-combine strategy for data analysis. Journal of Statistical Software 40, 1–29 (2011).

83 Wickham, H. ggplot2: elegant graphics for data analysis. (springer, 2016).

84 Slowikowski, K. et al. Package ggrepel. Automatically Position Non-Overlapping Text Labels with ‘ggplot2 (2018).

85 Brownrigg, R., Minka, T. & Deckmyn, A. maps: Draw Geographical Maps. R package version 3.3.0. (2018).

86 Becker, R., Wilks, A. & Brownrigg, R. mapdata: Extra map databases. R package version 2.3.0. (2018).

87 Gu, Z., Gu, L., Eils, R., Schlesner, M. & Brors, B. circlize implements and enhances circular visualization in R. Bioinformatics 30, 2811–2812 (2014).

88 Hope, R. M. ‘Rmisc: Ryan miscellaneous’. R package version 1.5.2 (2013).

89 Han, M. V. & Zmasek, C. M. phyloXML: XML for evolutionary biology and comparative genomics. BMC bioinformatics 10, 1–6 (2009).

90 Paradis, E. & Schliep, K. ape 5.0: an environment for modern phylogenetics and evolutionary analyses in R. Bioinformatics 35, 526–528 (2019).

