## Supplementary Figure 1 for "Polygenic adaptation from standing genetic variation allows rapid ecotype formation"

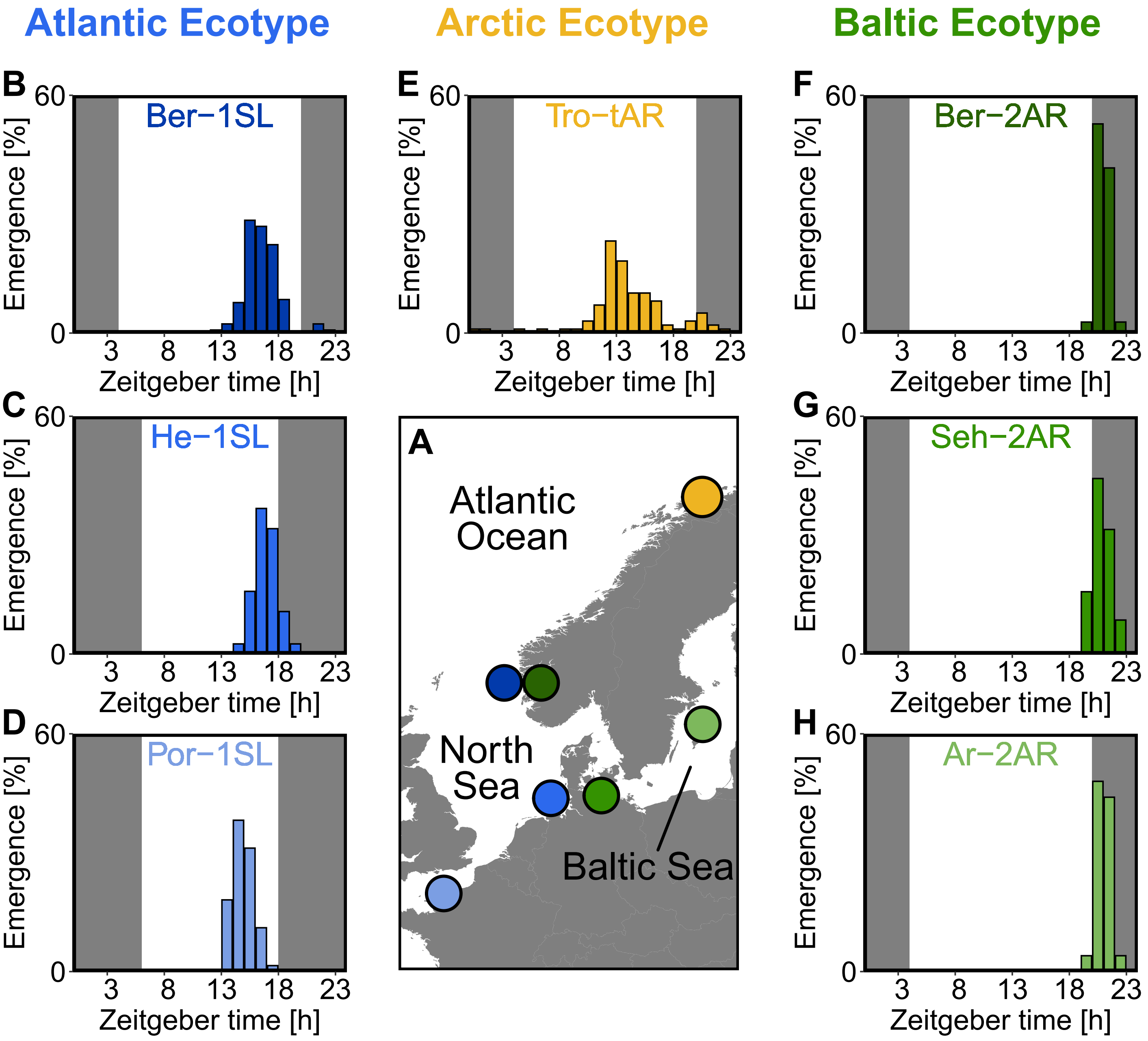


**Supplementary Figure 1 Circadian emergence rhythm of the studied *Clunio* strains under laboratory conditions.**

**(A)** Geographic locations. **(B-H)** Circadian emergence rhythms of laboratory strains. Dark shading indicates the dark phase, the middle of dark phase is defined as zeitgeber time 0. Data for Ber-1SL (B; n=130), Ber-2AR (F; n=36) and Tro-tAR (E; n=99) was recorded with a custom-made fraction collector in 1-h intervals under an artificial light cycle with 16 h of light and 8 h of darkness (LD 16:8). Data for Seh-2AR (G; n=70) and Ar-2AR (H; n=25) was recorded manually while performing crosses. Data of He-1SL (C) and Por-1SL (D) was taken from Neumann, 1966 and recorded under LD 12:12.
