## Supplementary Figure 2 for "Polygenic adaptation from standing genetic variation allows rapid ecotype formation"

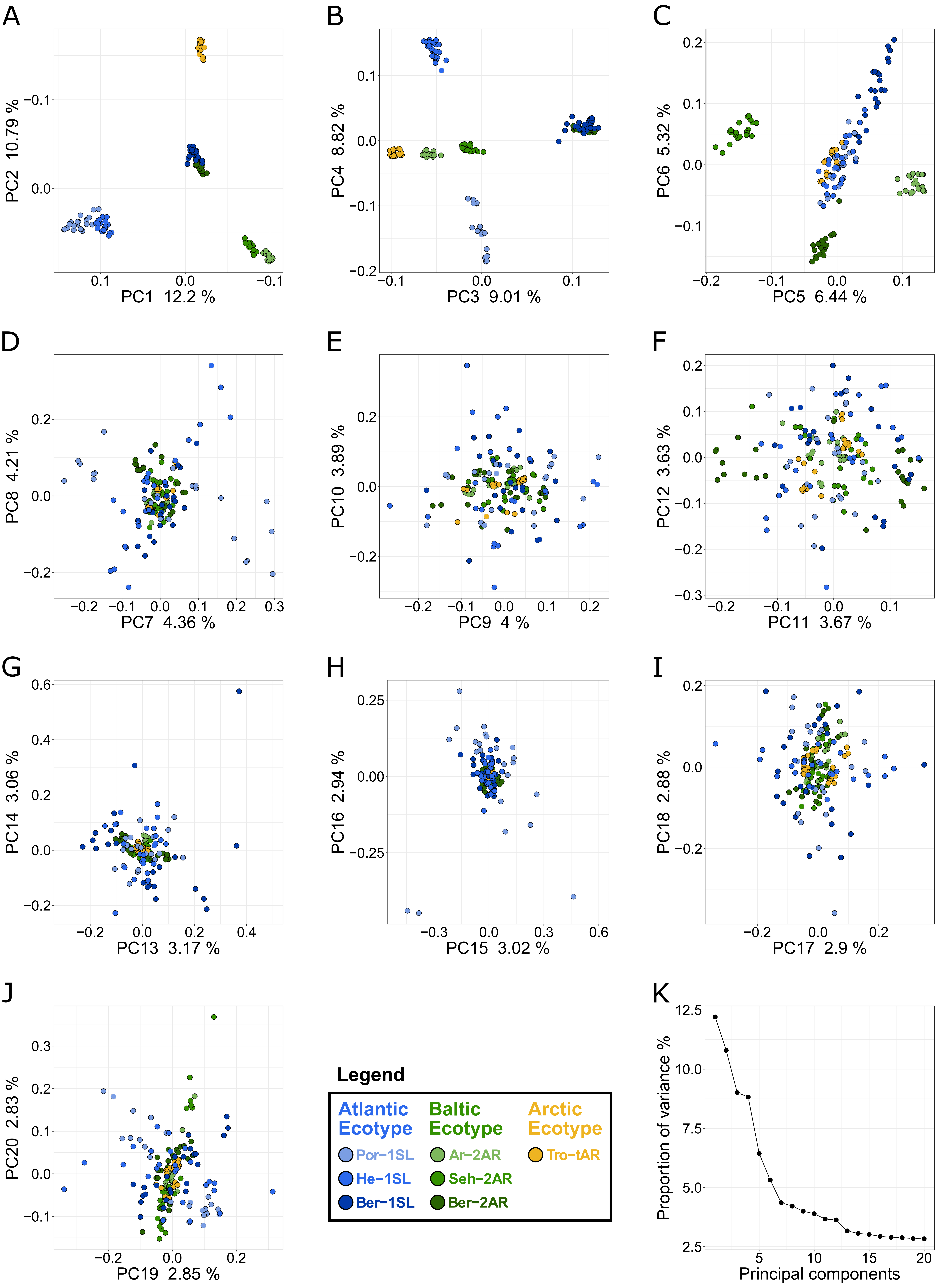


**Supplementary Figure 2 Principal component analysis (PCA) of all individuals for all seven populations.**

(**A-J**) The principal components (PCs) are plotted in pairs from 1-2 (A) to 19-20 (J). The eigenvalue as fraction of total variance is given on the respective axes. The individuals are color coded according to population (see legend). (**K**) Overview of the eigenvalue as fraction of total variance for all 20 PCs.
