## Supplementary Figure 3 for "Polygenic adaptation from standing genetic variation allows rapid ecotype formation"

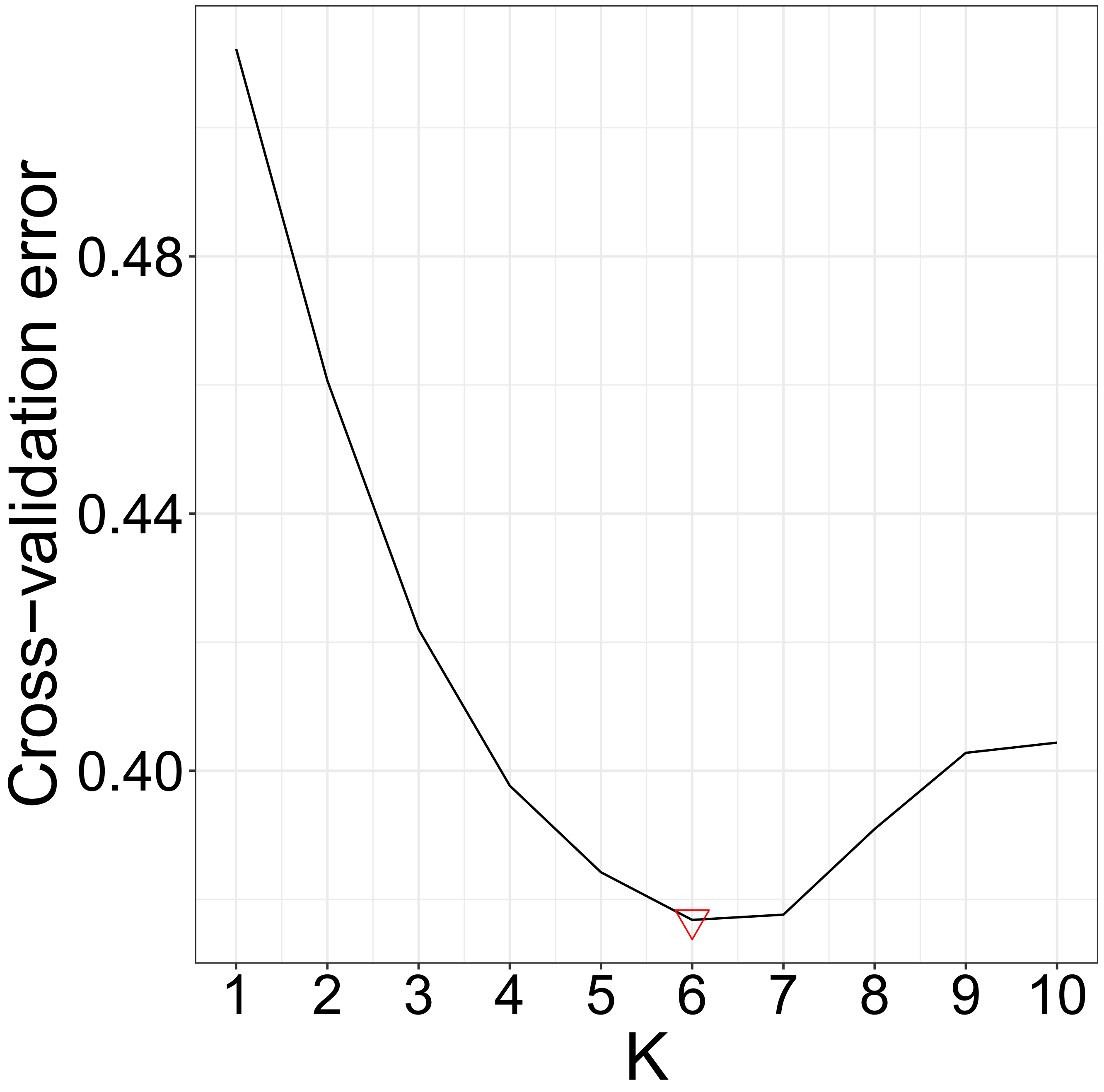


**Supplementary Figure 3 The results of the cross-validation test of the ADMIXTURE analysis.**

The optimal K is marked with a red downward directed triangle.
