## Supplementary Figure 4 for "Polygenic adaptation from standing genetic variation allows rapid ecotype formation"

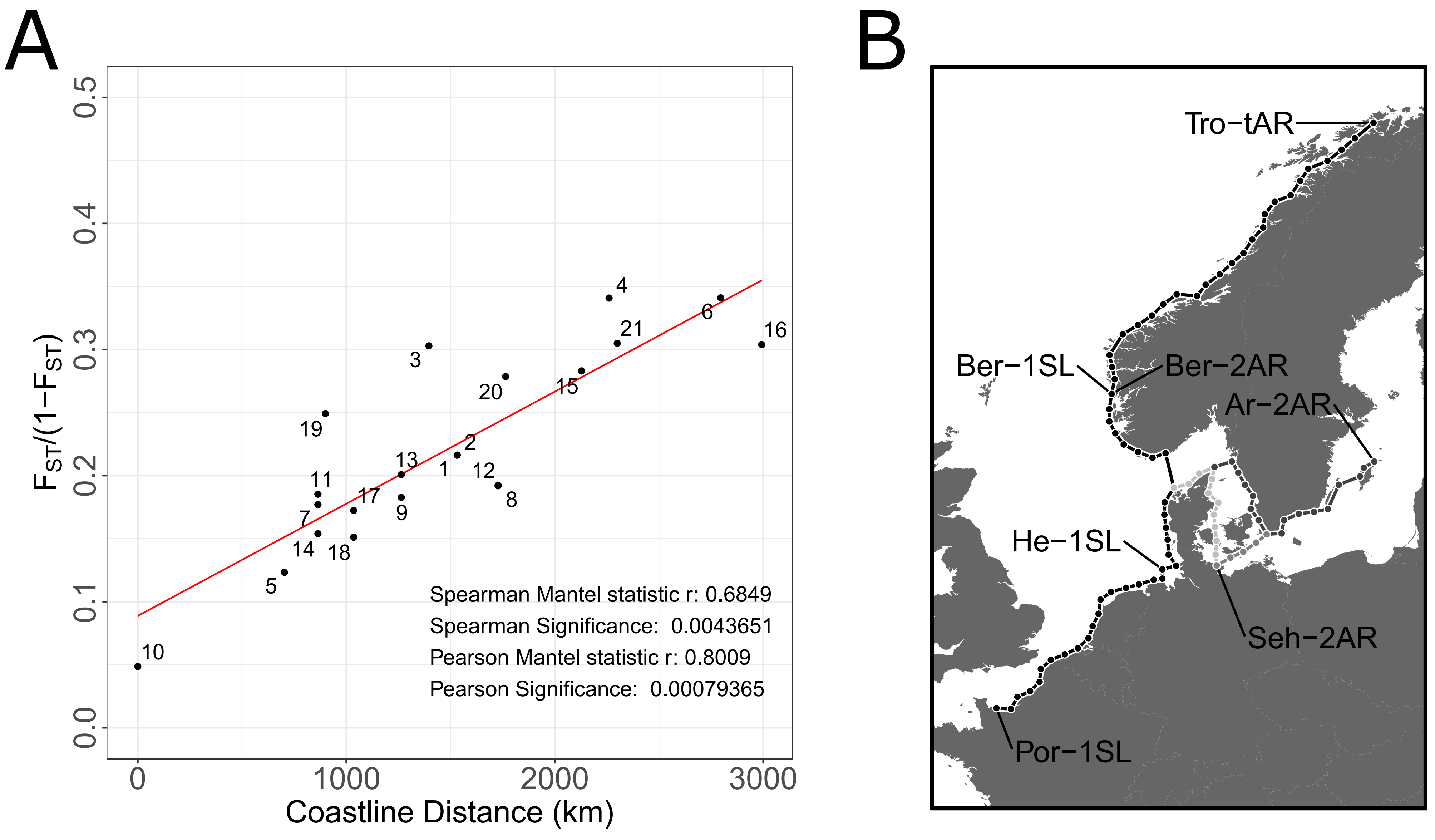


C

| Number | Compared populations | Estimated coastline distance (km) | $F_{ST}$ | $F_{ST}/(1-F_{ST})$ |
|---|---|---|---|---|
| 1 | Ar-2AR\|Ber-1SL | 1532,143 | 0.1509485 | 0.1777849 |
| 2 | Ar-2AR\|Ber-2AR | 1532,143 | 0.1509755 | 0.1778222 |
| 3 | Ar-2AR\|He-1SL | 1396,429 | 0.1886286 | 0.2324811 |
| 4 | Ar-2AR\|Por-1SL | 2260,715 | 0.202672 | 0.254189 |
| 5 | Ar-2AR\|Seh-2AR | 703,5714 | 0.09887547 | 0.1097245 |
| 6 | Ar-2AR\|Tro-tAR | 2796,429 | 0.2027245 | 0.2542715 |
| 7 | Ber-1SL\|He-1SL | 864,2857 | 0.1307079 | 0.1503613 |
| 8 | Ber-1SL\|Por-1SL | 1728,571 | 0.1386897 | 0.1610218 |
| 9 | Ber-1SL\|Tro-tAR | 1264,286 | 0.1338149 | 0.1544877 |
| 10 | Ber-2AR\|Ber-1SL | 0 | 0.04425415 | 0.04630327 |
| 11 | Ber-2AR\|He-1SL | 864,2857 | 0.1351794 | 0.1563091 |
| 12 | Ber-2AR\|Por-1SL | 1728,571 | 0.1389443 | 0.1613651 |
| 13 | Ber-2AR\|Tro-tAR | 1264,286 | 0.1432648 | 0.1672217 |
| 14 | He-1SL\|Por-1SL | 864,2857 | 0.1176835 | 0.1333802 |
| 15 | He-1SL\|Tro-tAR | 2128,572 | 0.1807271 | 0.2205945 |
| 16 | Por-1SL\|Tro-tAR | 2992,857 | 0.1890308 | 0.2330925 |
| 17 | Seh-2AR\|Ber-1SL | 1035,714 | 0.1281758 | 0.1470202 |
| 18 | Seh-2AR\|Ber-2AR | 1035,714 | 0.1160381 | 0.1312705 |
| 19 | Seh-2AR\|He-1SL | 900 | 0.1662999 | 0.1994721 |
| 20 | Seh-2AR\|Por-1SL | 1764,286 | 0.1788937 | 0.2178691 |
| 21 | Seh-2AR\|Tro-tAR | 2300 | 0.1894261 | 0.2336938 |

**Supplementary Figure 4 Mantel test for isolation by distance between the studied populations.**

**(A)** The average genomic differentiation is plotted against the estimated coastline distance on a pairwise level. Results of the Spearman and Pearson Mantel statistic are noted in the lower right corner of the graph. A trendline is drawn in red. **(B)** The coastline distance was hand-measured in 50 km intervalls. Distances between ceratin locations had different shortest distances, which are colored in different gray tones. **(C)** Every pairwise comparison is listed by number, the population names, the estimated coastline distance in km, their average $F_{ST}$ value and the $F_{ST}/(1-F_{ST})$ values.
