## Supplementary Figure 5 for "Polygenic adaptation from standing genetic variation allows rapid ecotype formation"

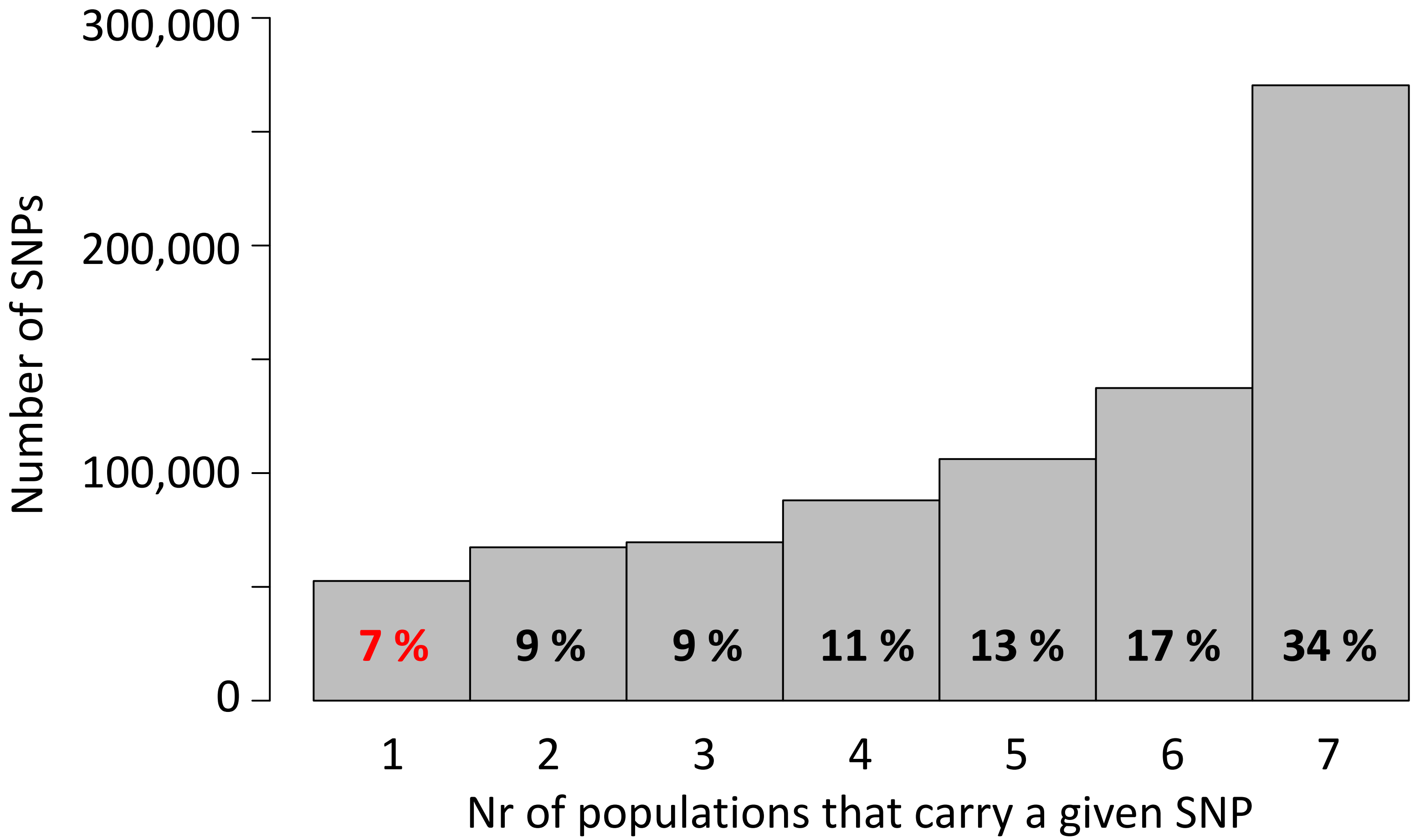


**Supplementary Figure 5 Detected single nucleotide polymorphisms (SNPs) are largely shared between the seven populations.**

Of a total of 792,032 SNPs, 34% are found in all seven populations, whereas only 7% are private to one of the seven populations.
