## Supplementary Figure 6 for "Polygenic adaptation from standing genetic variation allows rapid ecotype formation"

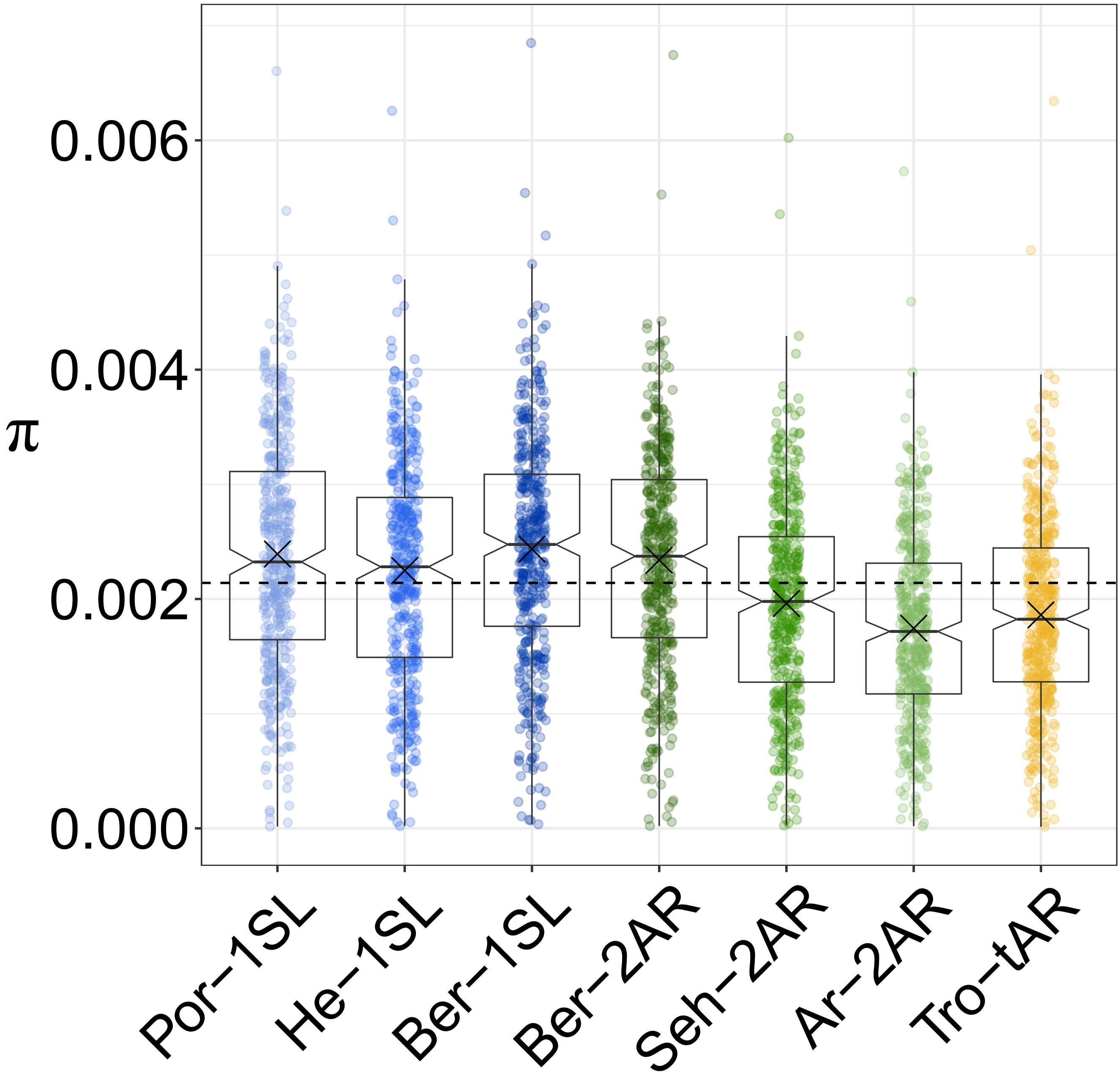


**Supplementary Figure 6 Nucleotide diversity π per population in 200 kb non-overlapping windows across the genome.**

Populations are colour coded as in Figure 1A and all individuals per population were used for the analysis. The boxplots show the median, 25th and 75th quantile, data maximum and minimum, and the outliers. The arithmetic mean is marked per boxplot as 'X' and the overall arithmetic mean as dashed line.
