## Supplementary Figure 7 for "Polygenic adaptation from standing genetic variation allows rapid ecotype formation"

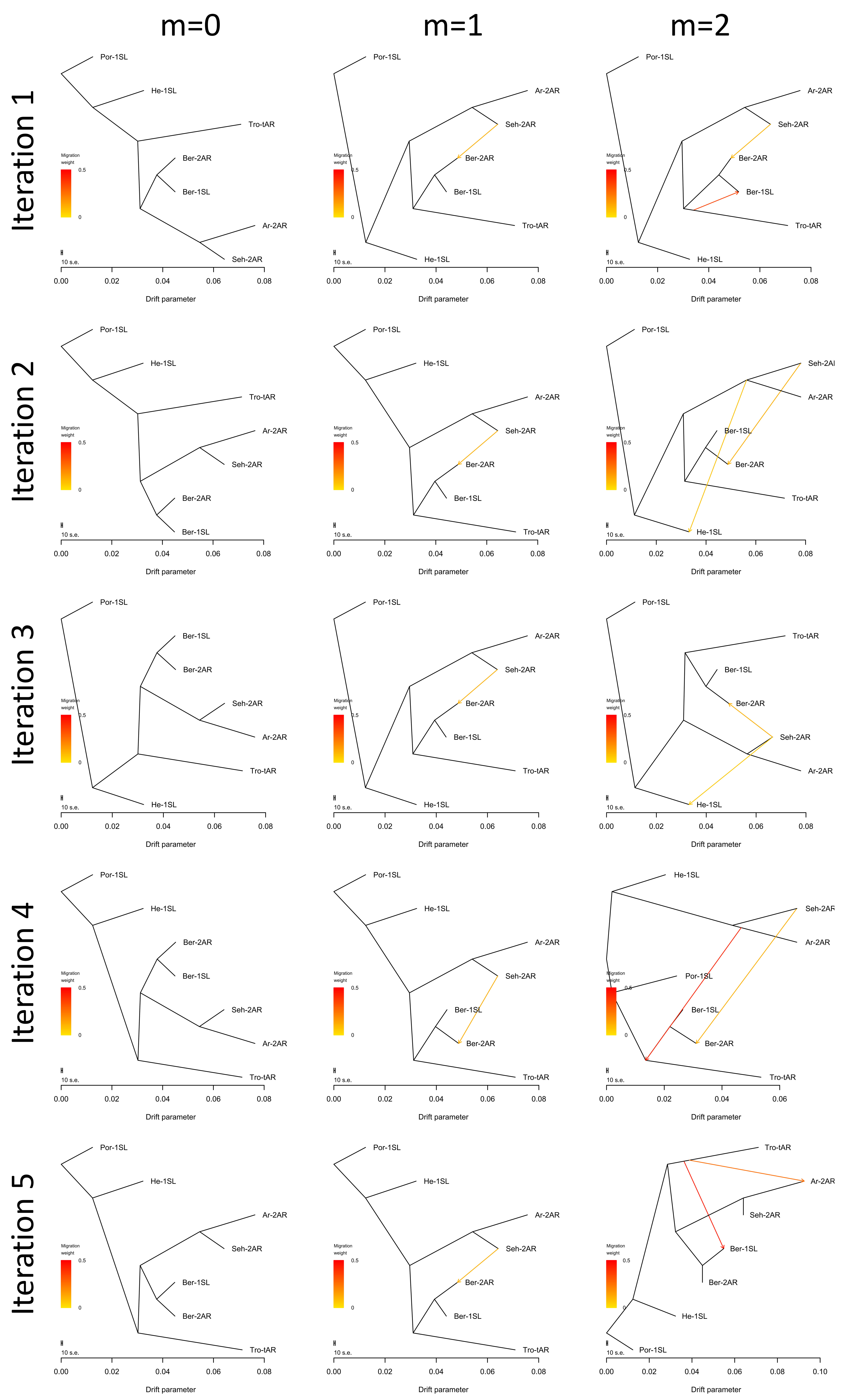


**Supplementary Figure 7 Treemix analysis for introgression events.**

Treemix was run for no, one or two migration events with five iterations each. Analysis with zero migration events yields the same structure of historic population splits in all five iterations, congruent with the genome-wide consensus phylogeny (Supplementary Fig. 9). With one migration event, the algorithm stably detects introgression from Seh-2AR into Ber-2AR. With two migration events, introgression from Seh-2AR into Ber-2AR is usually still detected (4 out of 5 iterations), but the second introgression event is random, suggesting there is no reliably detectable second introgression event.
