## Supplementary Figure 8 for "Polygenic adaptation from standing genetic variation allows rapid ecotype formation"

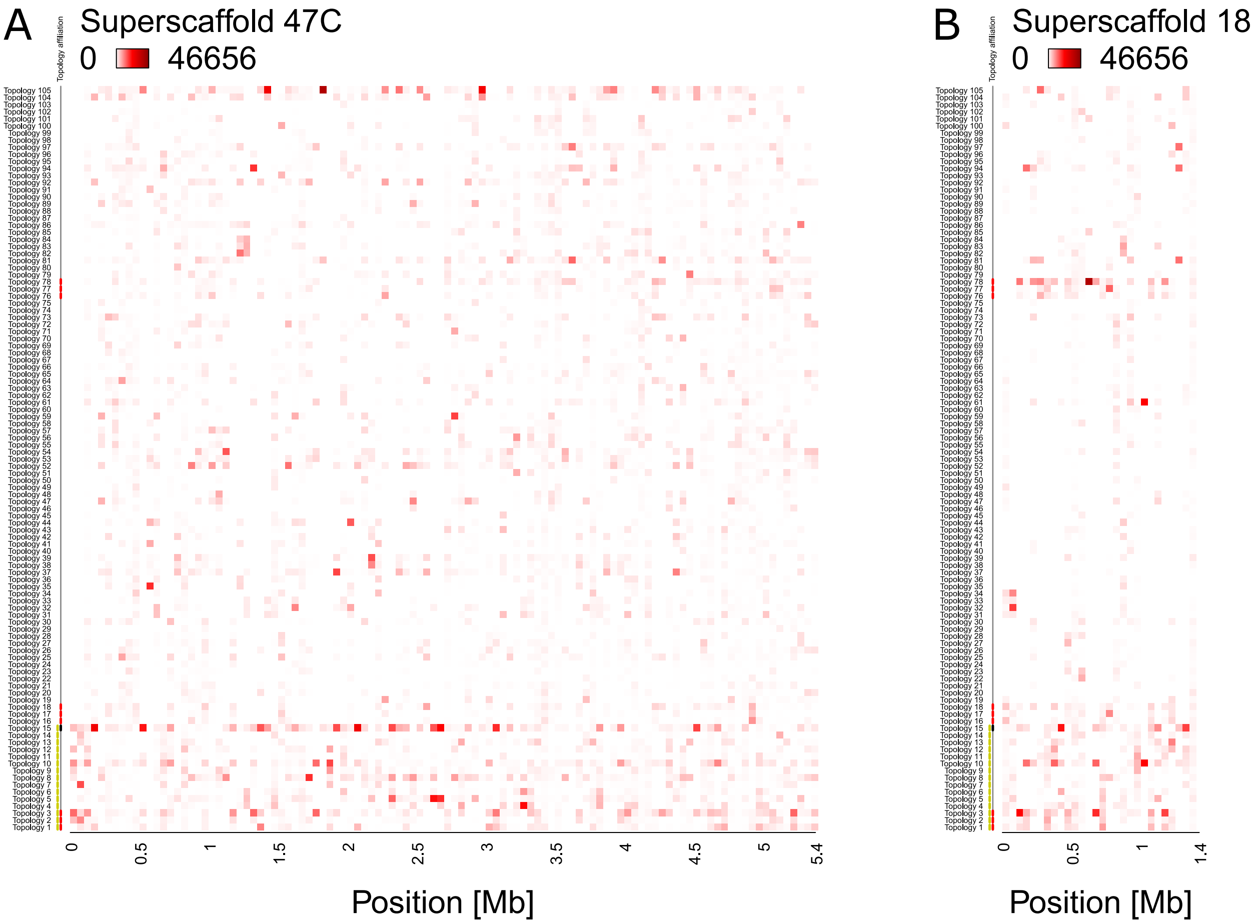


**Supplementary Figure 8 Incomplete lineage sorting, as illustrated by the relative support for 105 population topologies in 50 kb windows across superscaffolds 47C and 18.**

The heatmap shows the assigned topology weight for each topology per window ranging from 0 (white, no support) to 46,656 (red, support by all trees). The supported topologies change quickly along the chromosome. In almost all windows, several topologies are supported, which implies that individuals do not cluster according to population, highlighting incomplete lineage sorting. Topology 15 (black mark on the left) is the whole-genome consensus population topology. Topologies marked in red are separating populations according to ecotypes. Topologies marked in yellow are consistent with introgression. **(A)** Superscaffold47C – the longest scaffold in the reference genome – gives an impression of the genomic background, with highest overall support for topology 15. **(B)** Superscaffold 18, from the middle of chromosome 1, is strongly enriched for ecotype-associated topologies (compare Figure 3).
