## Supplementary Figure 9 for "Polygenic adaptation from standing genetic variation allows rapid ecotype formation"

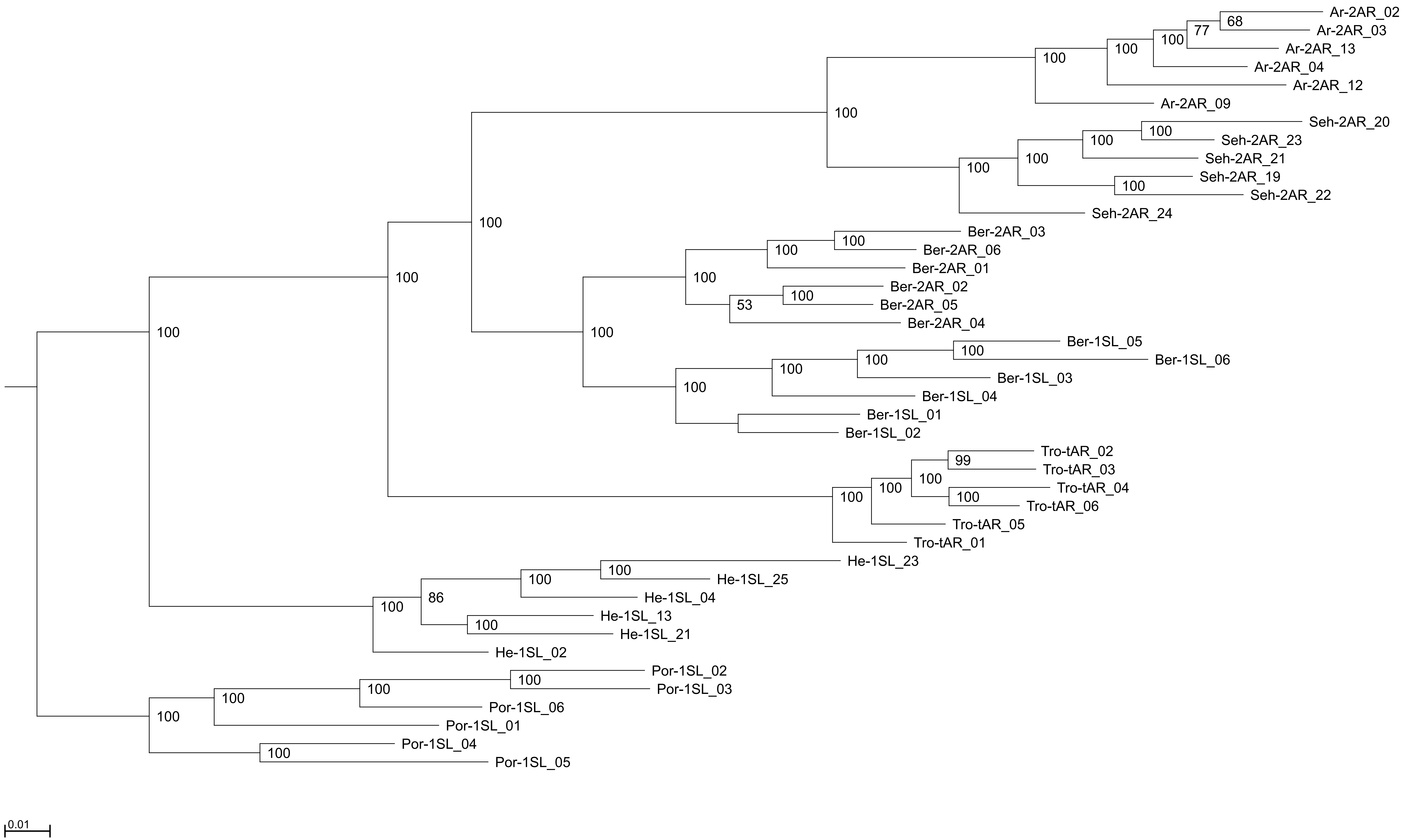


**Supplementary Figure 9 Genome-wide consensus genealogy for six individuals from each of the seven populations.**

The same six individuals from each population were also used in the TWISST topology weighting analysis. The genealogy is based on the whole genome SNP sequences, composed of 792,032 positions. The SNPs were extracted from the VCF using the vcf2phylip.py v.2.3 script. The genealogy was calculated with IQ-TREE v. 1.6.12 and graphically edited with Archaeopteryx v.0.968.
