## Supplementary Figure 10 for "Polygenic adaptation from standing genetic variation allows rapid ecotype formation"

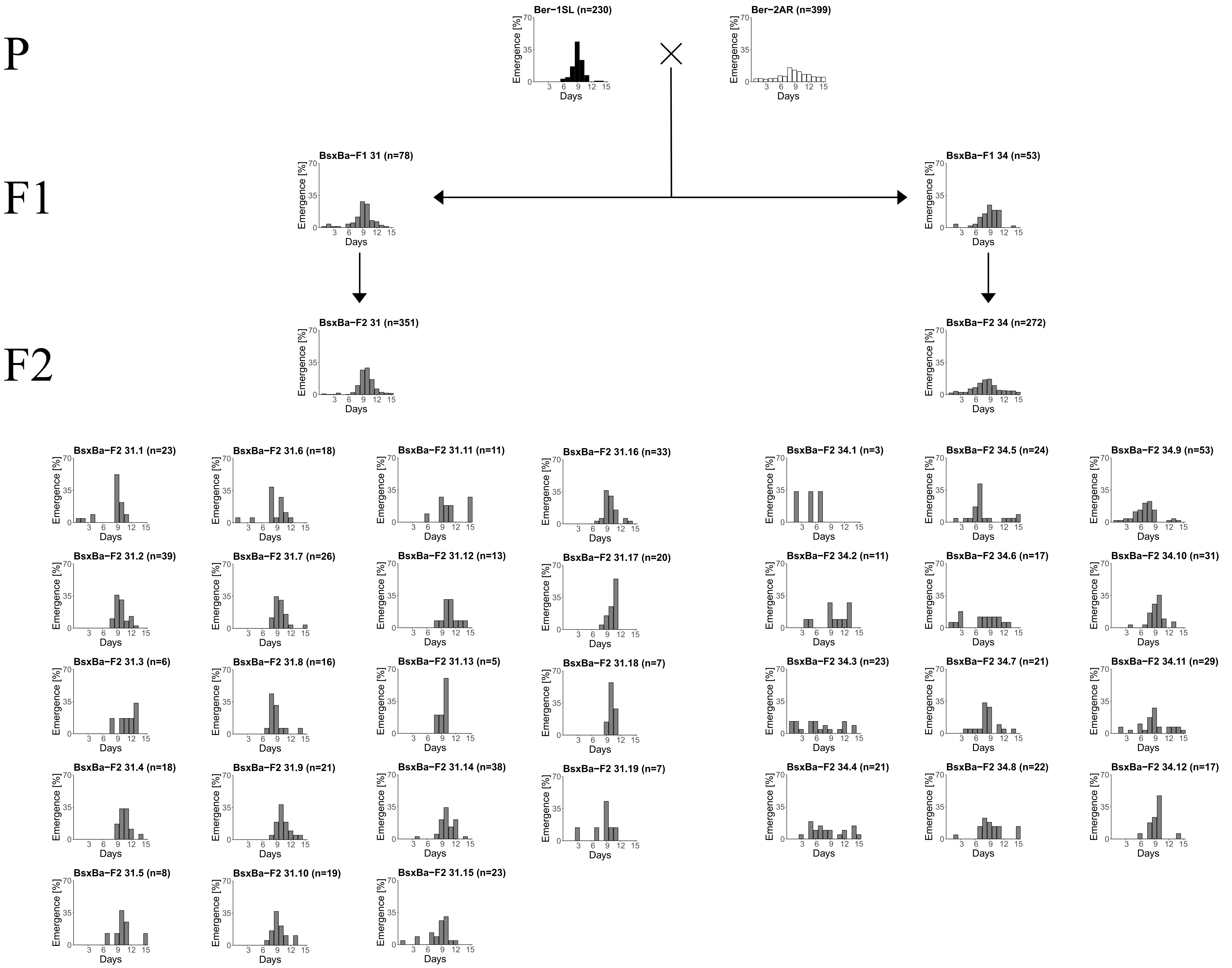


**Supplementary Figure 10 Lunar emergence patterns of crosses between the Ber-1SL and Ber-2AR strains.**

Single pair crosses were set up between the Ber-1SL and Ber-2AR strains, resulting in several F1 families, two of which are shown here (F1-31 and F1-34). F2 families were obtained by letting the siblings within each F1 family mate freely with each other. Thus, from each F1 we obtained several F2 families, which all go back to a single pair of parents. F1-31 and F1-34 differ in the degree of lunar rhythmicity, as do F2-31 and F2-34. When plotting the F2 families individually, it becomes apparent that in F2-31 all families are quite rhythmic. In contrast, in F2-34 there are highly rhythmic families (e.g. F2-34.10) and completely arrhythmic families (e.g. F2-34.3 and F2-34.4), suggesting that this F2 generation is segregating for rhythmicity alleles. The observation suggests that in the cross leading to F1-31/F2-31, the Ber-2AR parent carried a considerable fraction of rhythmic alleles (basically “Ber-1SL” alleles), which rendered the resulting F1 and F2 generations largely rhythmic. The Ber-2AR parent for the F1-34/F2-34 cross seemed to carry largely arrhythmic alleles, as expected, which then segregate in the F2. Overall, we conclude that the Ber-2AR strain seems to carry a certain fraction of lunar rhythmic alleles. The segregation of lunar rhythmicity not only within but also between F2 families suggests a heterogeneous polygenic architecture of the trait.
