## Supplementary Figure 11 for "Polygenic adaptation from standing genetic variation allows rapid ecotype formation"

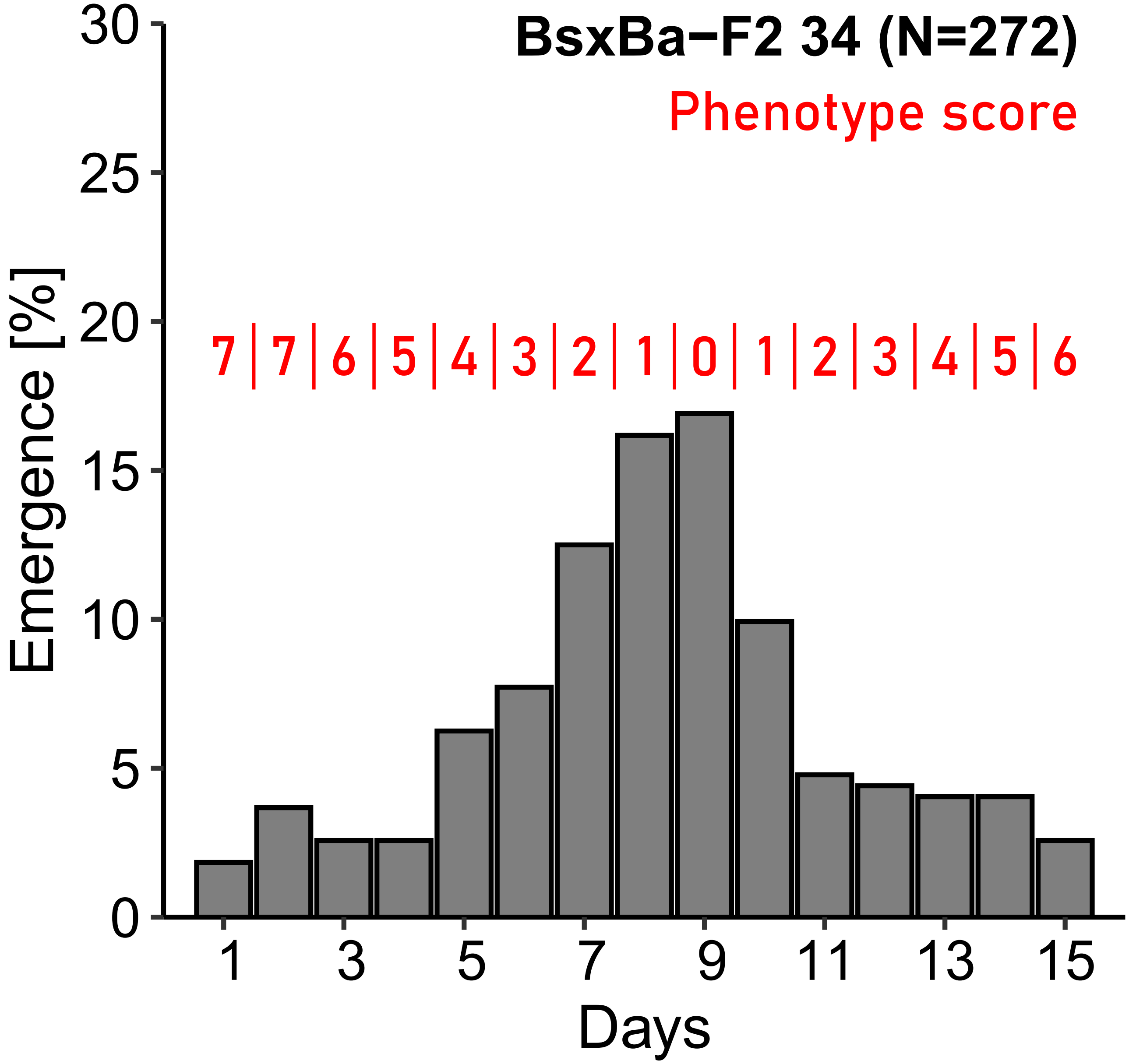


**Supplementary Figure 11 Phenotyping of the F2 individuals based on their emergence day.**

Lunar rhythmicity in the individuals was scored based on their emergence day, in relation to the Ber-1SL emergence distribution peak on day 9 of the 15-day turbulence cycle.
