## Supplementary Figure 12 for "Polygenic adaptation from standing genetic variation allows rapid ecotype formation"

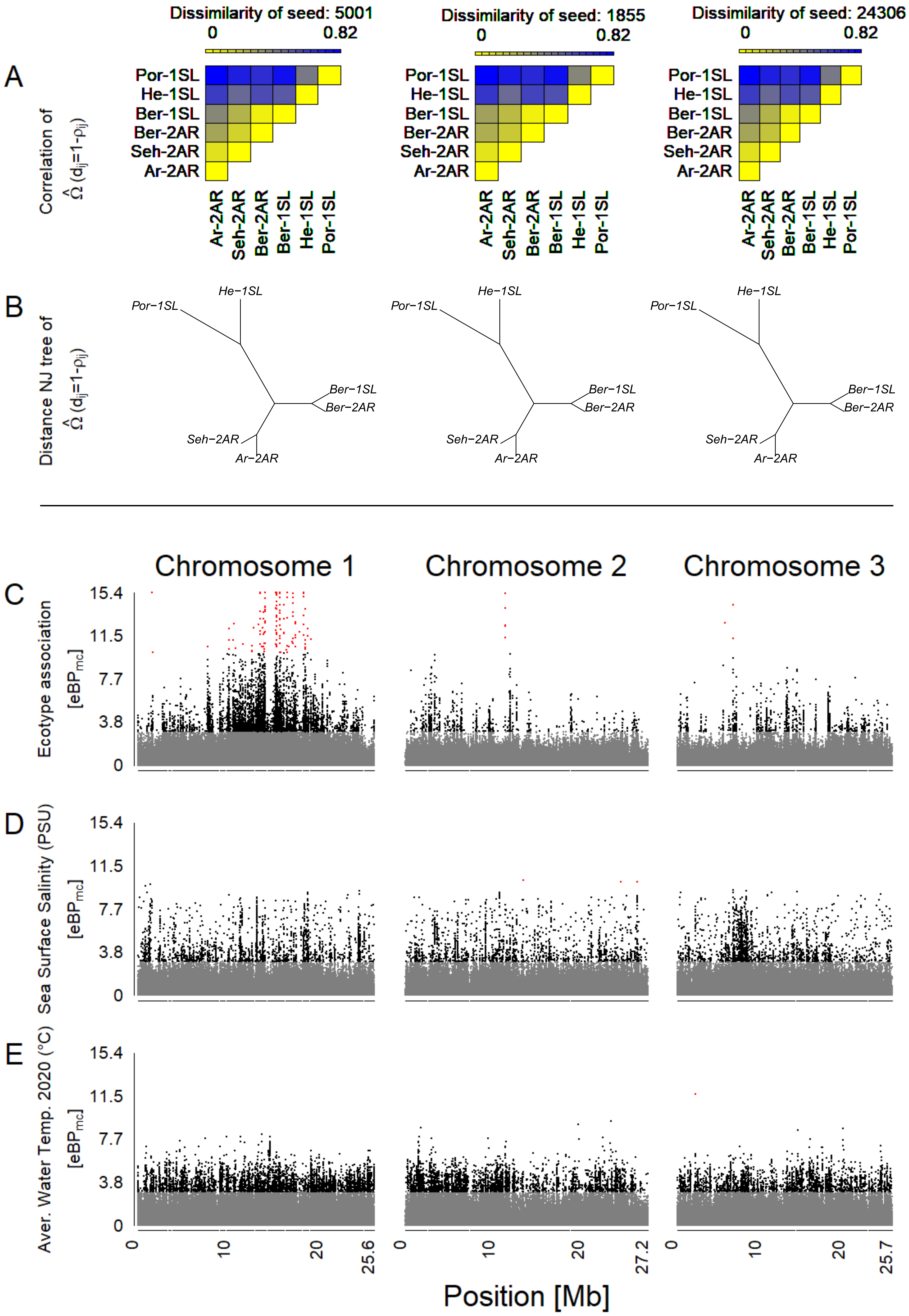


**Supplementary Figure 12 An overview of outlier and association analysis for ecotype (C), sea surface salinity (D) and water temperature (E).**

(**A**) Before assessment of outlier loci and association, the dataset was corrected for population structure based on an Omega matrix, which is here visualized as a dissimilarity matrix. Three independent runs (seeds 5001, 1855 and 24306) converge. (**B**) Omega matrices can also be visualized as Neighbor Joining trees. (**C-E**) Association of genetic variants with ecotype (C), sea surface salinity (D) and water temperature (E) is assessed via the $eBP_{mc}$ score and plotted along the three chromosomes, color-coded in grey ($eBP_{mc} < 3$), black ($3 < eBP_{mc} < 10$) and red ($eBP_{mc} > 10$).
