## Supplementary Figure 13 for "Polygenic adaptation from standing genetic variation allows rapid ecotype formation"

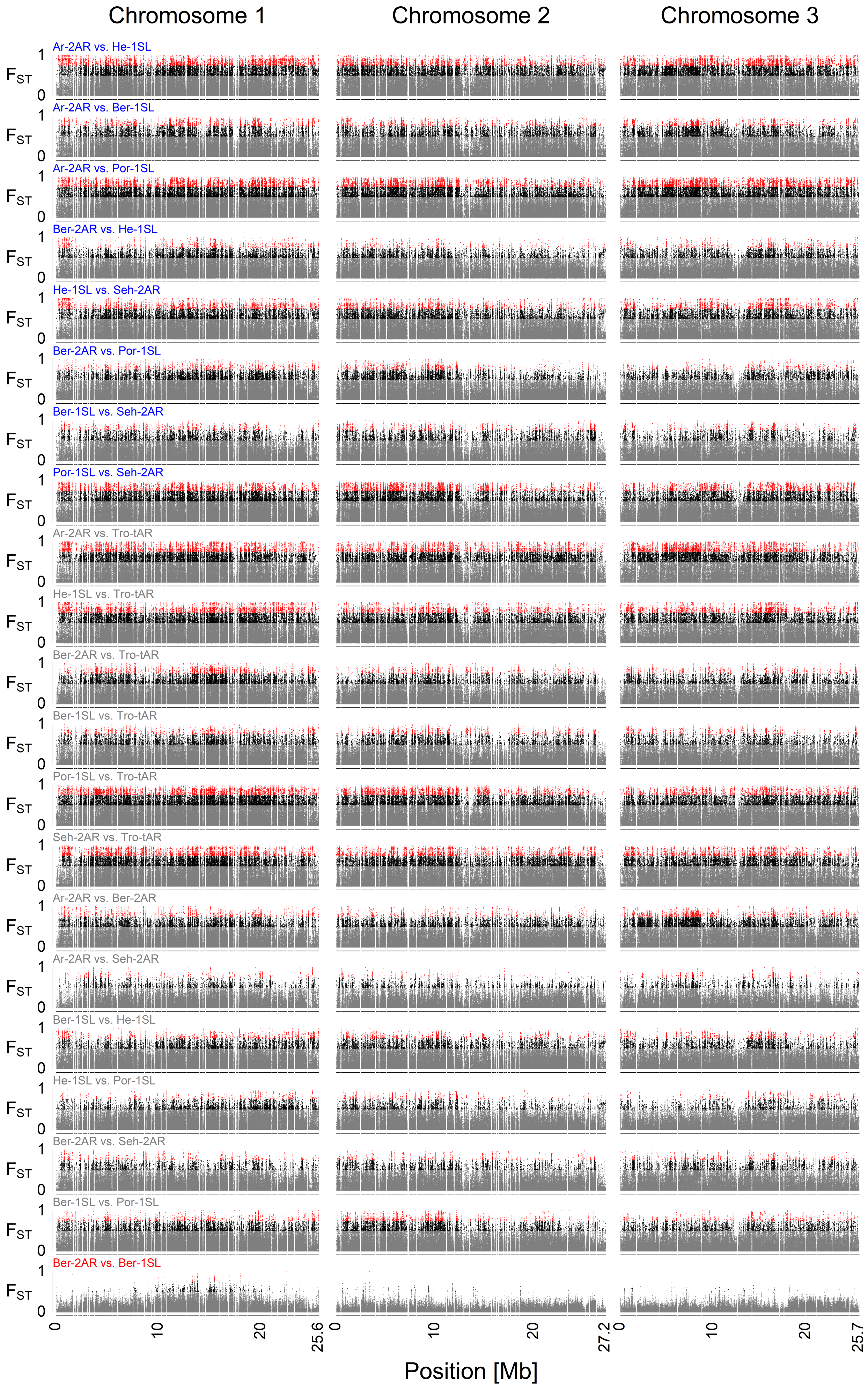


**Supplementary Figure 13 Pairwise $F_{ST}$ comparisons between all seven populations.**

The specific populations of the comparison are written above each plot, color-coded for specific comparisons. The comparison between the sympatric Ber-2AR and Ber-1SL ecotypes is plotted in red. Other comparisons of *Atlantic* vs. *Baltic ecotype* populations are plotted in blue. All other comparisons are plotted in grey. The $F_{ST}$ values are colour coded in grey ($F_{ST} < 0.5$), black ($0.5 < F_{ST} < 0.75$) and red ($F_{ST} > 0.75$). The sympatric ecotypes from Bergen are much less differentiated than all other populations.
