## Supplementary Figure 14 for "Polygenic adaptation from standing genetic variation allows rapid ecotype formation"

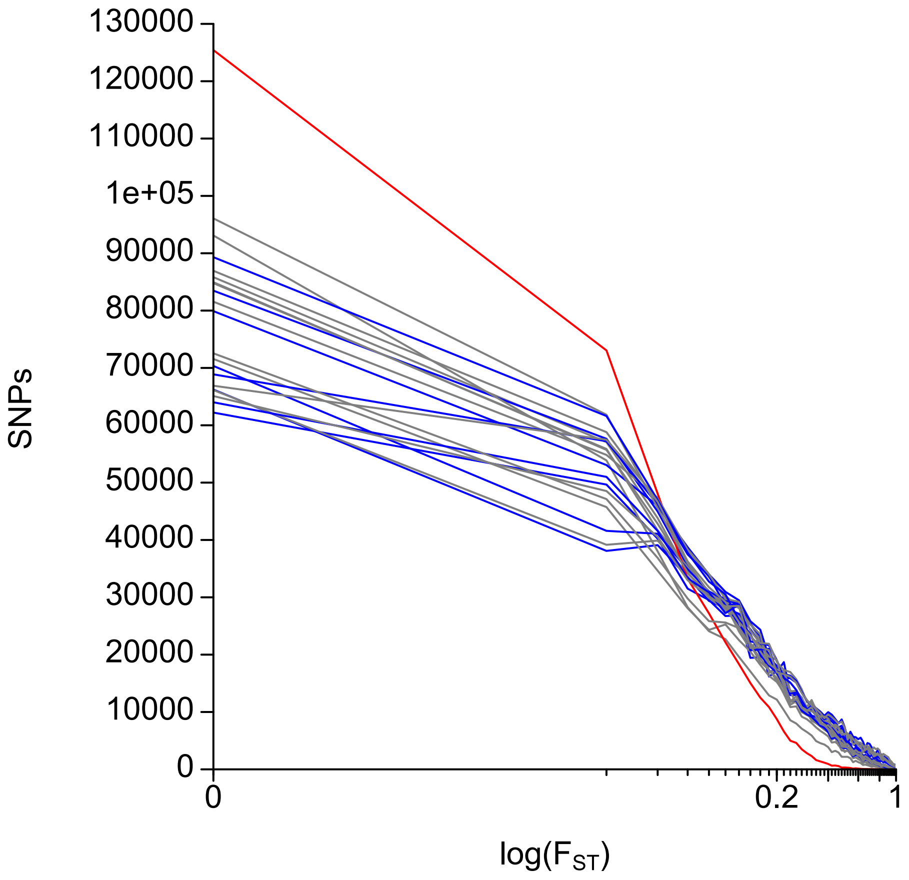


**Supplementary Figure 14 Distribution of $F_{ST}$ values in all pairwise population comparisons.**

$F_{ST}$ values for 792,032 SNPs were categorized in 0.02 bins. The comparison between the sympatric Ber-2AR and Ber-1SL ecotypes is plotted in red. Other comparisons of *Atlantic* vs. *Baltic ecotype* populations are plotted in blue. All other comparisons are plotted in grey.
