## Supplementary Figure 15 for "Polygenic adaptation from standing genetic variation allows rapid ecotype formation"

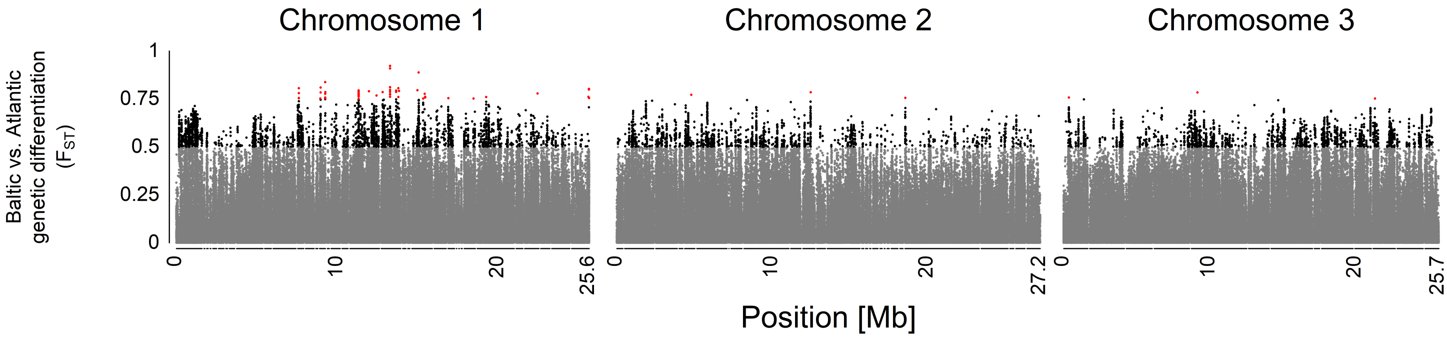


**Supplementary Figure 15 Genetic differentiation ($F_{ST}$) between Atlantic and Baltic ecotype.**

*Atlantic ecotype* populations (Por-1SL, He-1SL, Ber-1SL) and *Baltic ecotype* populations (Ar-2AR, Seh-2AR, Ber-2AR) were grouped. The $F_{ST}$ values are color-coded in grey ($F_{ST} < 0.5$), black ($0.5 < F_{ST} < 0.75$) and red ($F_{ST} > 0.75$).
