## Supplementary Figure 16 for "Polygenic adaptation from standing genetic variation allows rapid ecotype formation"

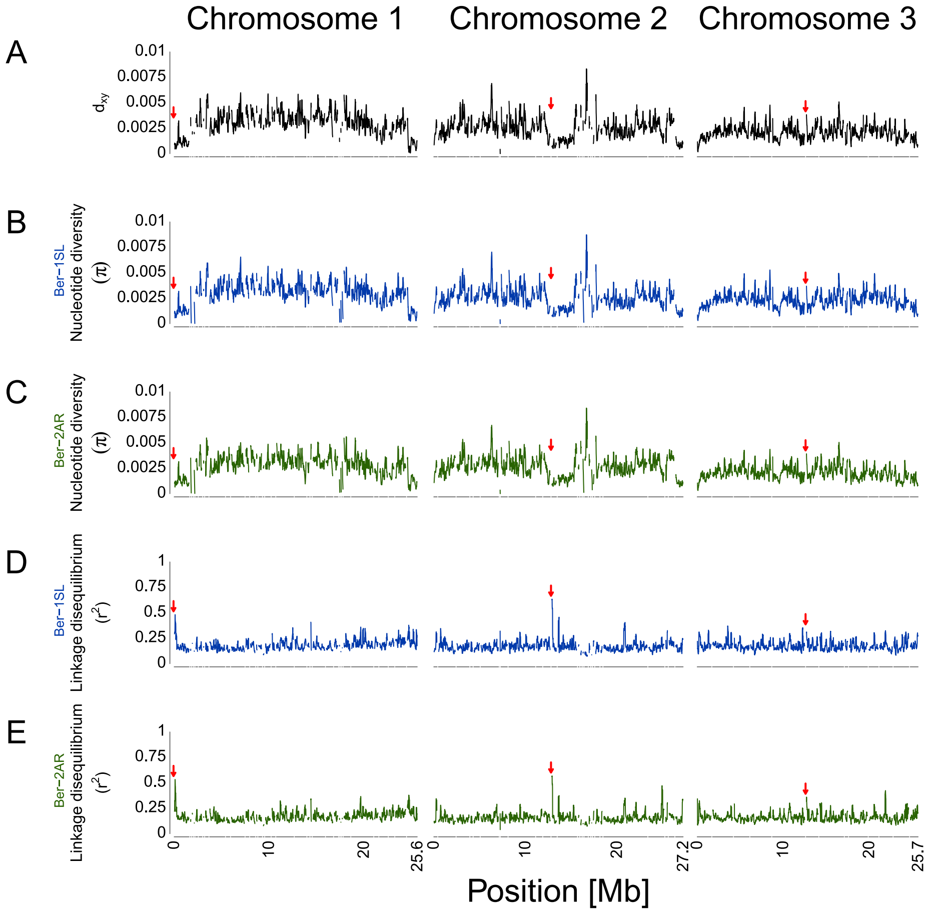


**Supplementary Figure 16 Genetic divergence ($d_{xy}$), nucleotide diversity ($\pi$) and short-range linkage disequilibrium ($r^2$) for the Ber-2AR vs. Ber-1SL comparison.**

Summary statistics are based on 792,032 SNPs and were calculated in 100 kb sliding windows with 10 kb steps. (**A**) Genetic divergence ($d_{xy}$) based on allele frequencies. (**B**, **C**) Nucleotide diversity $\pi$. (**D**, **E**) Linkage disequilibrium measured as $r^2$. Ber-1SL and Ber-2AR are given in their respective population colors. The approximate position of the centromeres are marked on each chromosome with red arrows. The position is based on the data provided by Kaiser et al., 2016.
