## Supplementary Figure 17 for "Polygenic adaptation from standing genetic variation allows rapid ecotype formation"

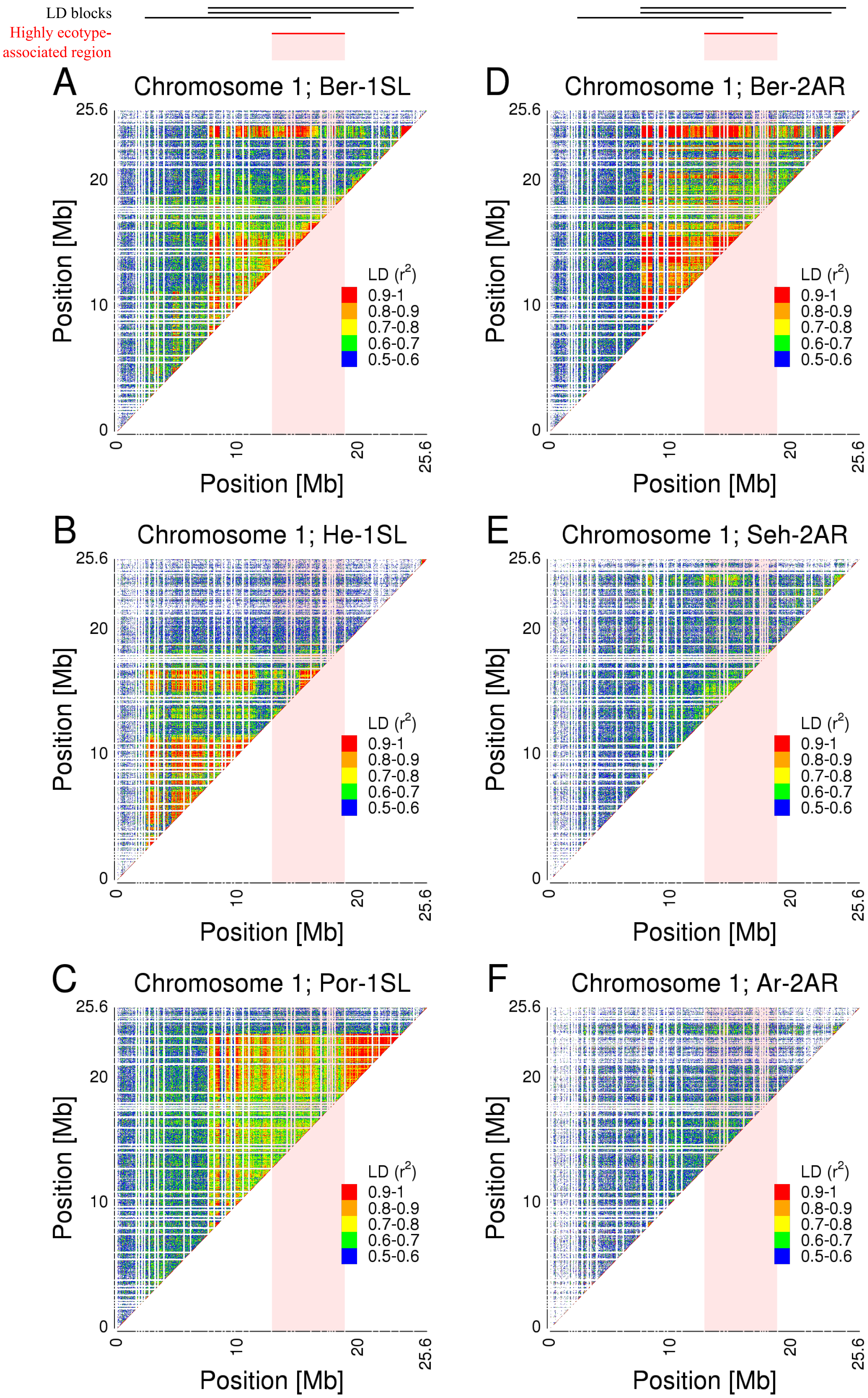


**Supplementary Figure 17 Pairwise linkage disequilibrium (LD) across chromosome 1 of all Atlantic and Baltic ecotype populations.**

Pairwise LD was calculated as '--geno-r2' in VCFtools with a '--maf 0.2' filter and a LD threshold of 0.5 $r^2$. The $r^2$ values are color coded from 0.5 (blue) to 1 (red) in 0.1 steps. The panels show the chromosome 1 $r^2$ values for each population (A: Ber-1SL, B: He-1SL, C: Por-1SL, D: Ber-2AR, E: Seh-2AR, F: Ar-2AR).
