## Supplementary Figure 18 for "Polygenic adaptation from standing genetic variation allows rapid ecotype formation"

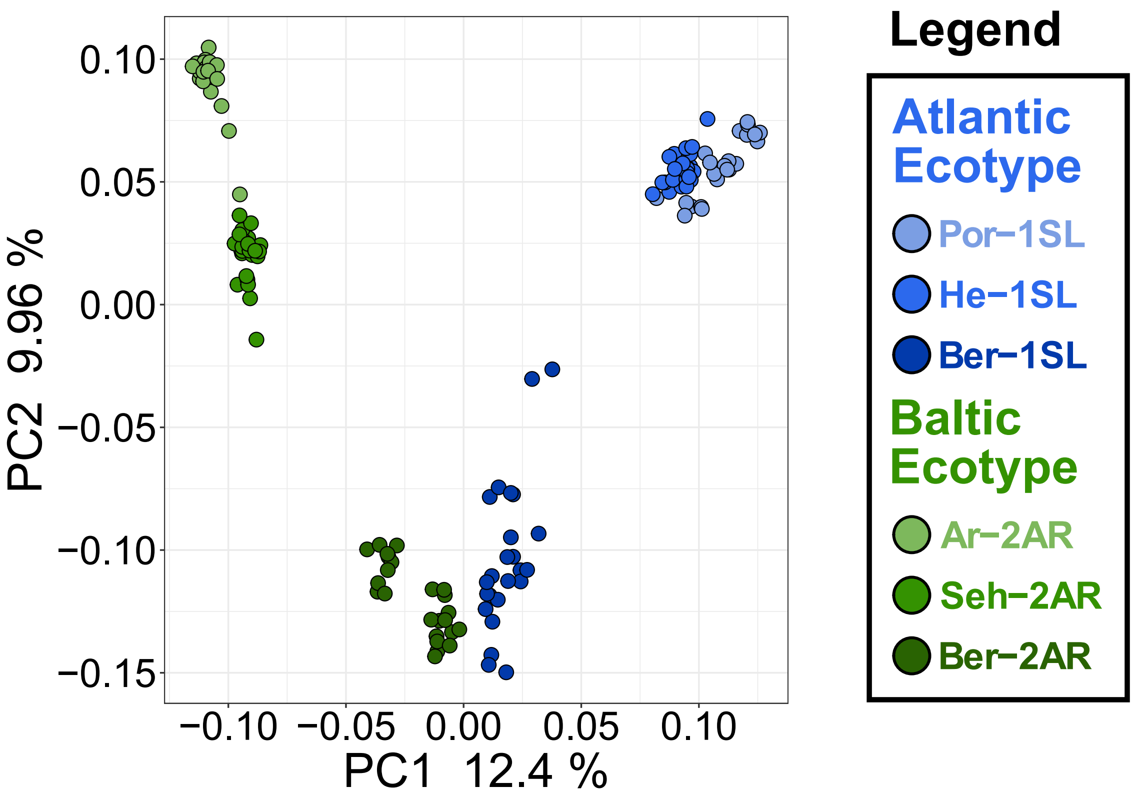


**Supplementary Figure 18 Principal component analysis (PCA) of the Atlantic and Baltic ecotypes for the highly ecotype-assocated region on chromosome 1.**

The highly ecotype-associated region on chromosome 1 ranges from superscaffolds 18 to 42. PCA for this region separates the ecotypes well, but does not show any patterns characteristic for a polymorphic structural variant (SV) separating the ecotypes. Only within the Por-1SL strain individuals cluster into three groups, suggestive of an overlapping polymorphic SV within that strain.
