## Supplementary Figure 19 for "Polygenic adaptation from standing genetic variation allows rapid ecotype formation"

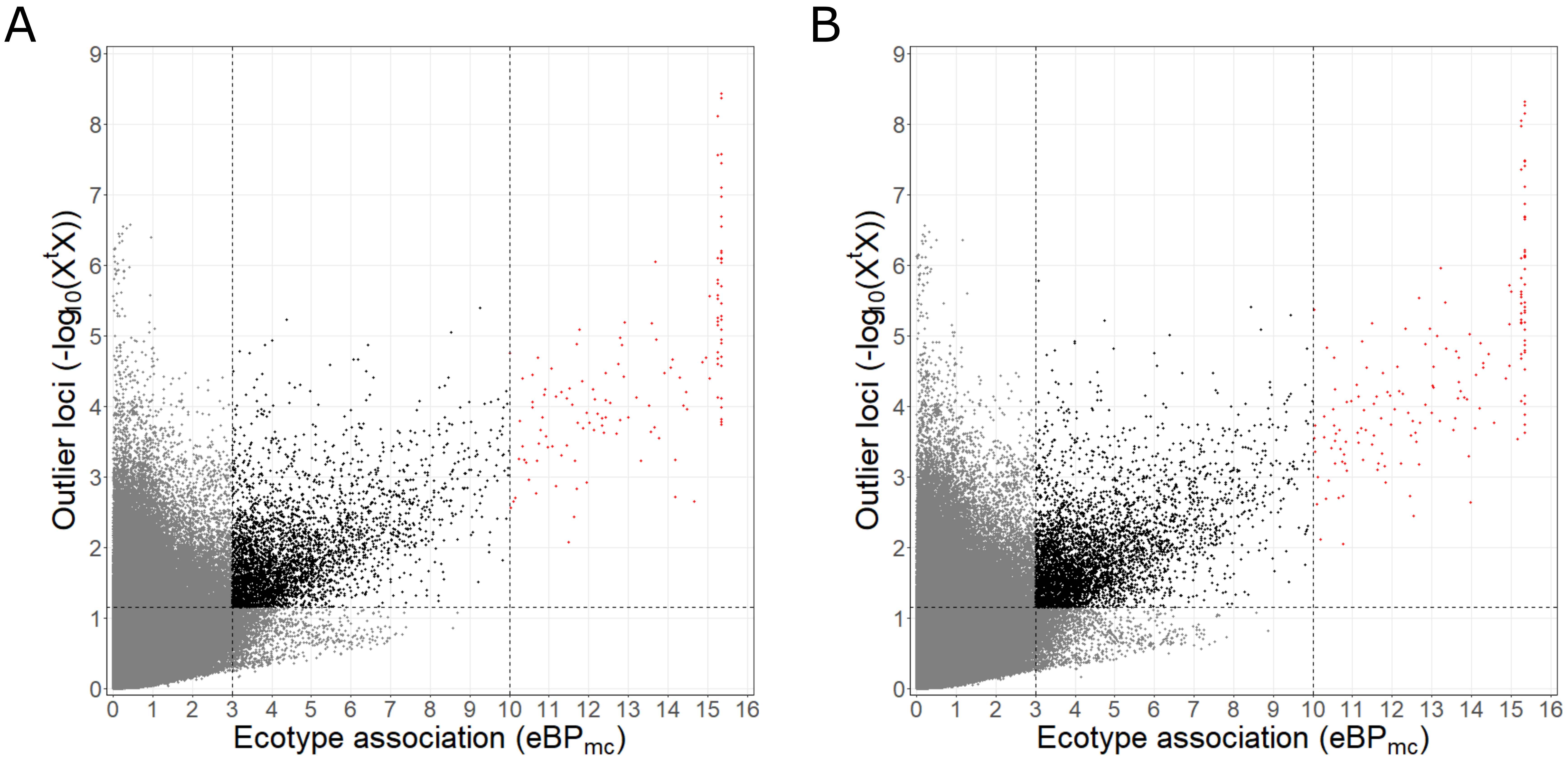


**Supplementary Figure 19 Selection of ecotype associated genetic variants based on $X^tX$ and $eBP_{mc}$ values.**

**(A)** Ecotype-associated SNPs were selected based on an $X^tX$ threshold of 1.152094 (as obtained from subsampling) and $eBP_{mc}$ thresholds of 3 or 10 respectively (corresponding to p-values for association of $10^{-3}$ or $10^{-10}$). SNPs above the $X^tX$ threshold and with $eBP_{mc} > 3$ are colored black, those with $eBP_{mc} > 10$ are colored in red. **(B)** Ecotype-associated variants (SNPs and indels) were selected based on an $X^tX$ threshold of 1.148764 and the same $eBP_{mc}$ thresholds.
