## Supplementary Figure 20 for "Polygenic adaptation from standing genetic variation allows rapid ecotype formation"

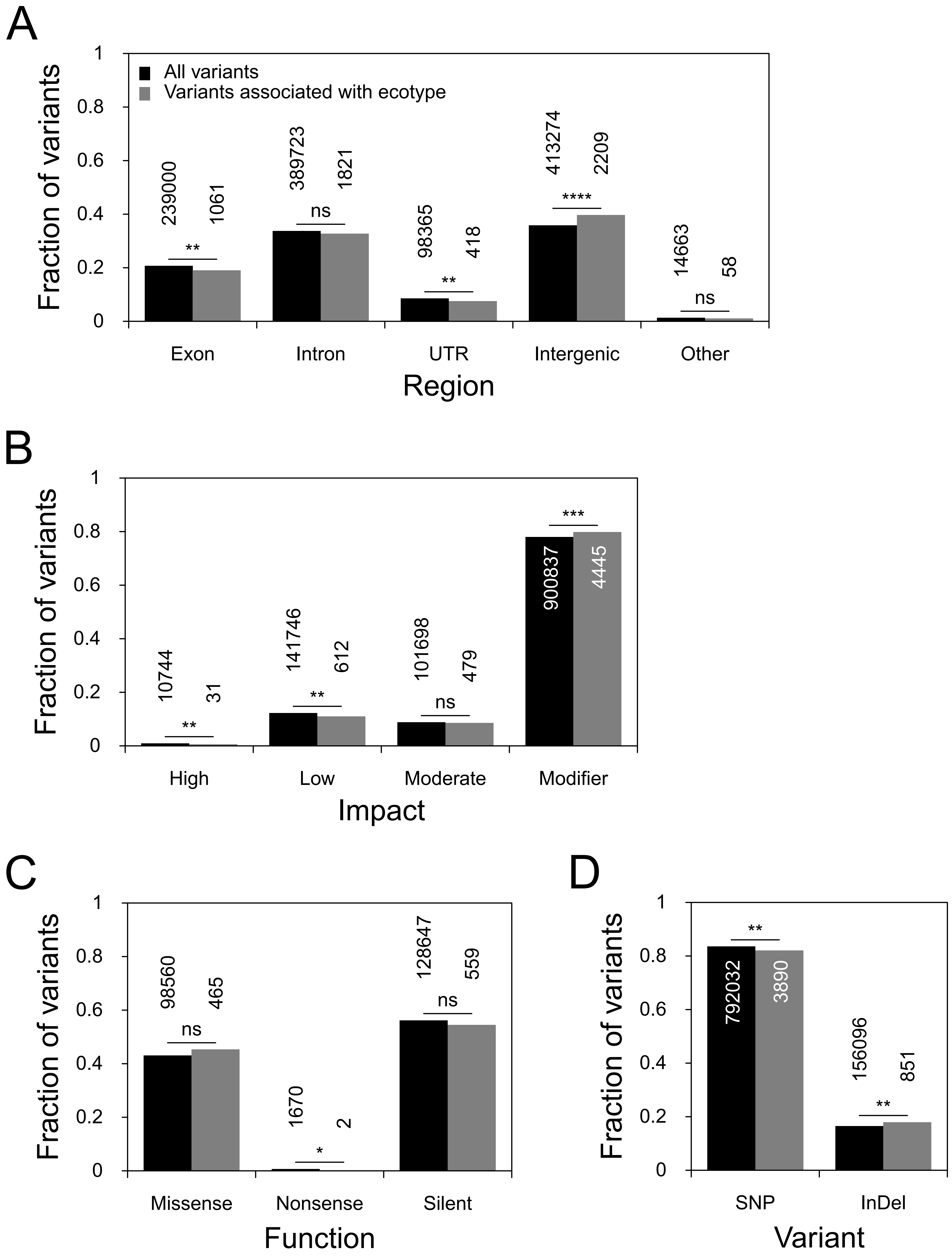


**Supplementary Figure 20 The effects of ecotype-associated genetic variants on genes, as compared to the genome-wide set of genetic variants.**

SnpEff assesses genetic variants for their position relative to annotated gene models and infers the effect that these variants have on the genes. We compared the locations and effects of all variants in the dataset (n=948,128; black bars) to those of ecotype-associated variants (n=4,741; grey bars) and tested for significant differences with Fisher's exact test. The absolute numbers for each class of variants are given above or inside the bar. **(A)** The location of genetic variants relative to gene models. **(B)** The estimated impact of these variants on the gene models. **(C)** The effect of variants that are found in coding regions. **(D)** The proportions of SNPs vs. indels. p-value symbols: **** - 0-0.0001; *** - 0.0001-0.001; ** - 0.001-0.01; * - 0.01-0.05; ns - 0.05-1.
