## Supplementary Figure 21 for "Polygenic adaptation from standing genetic variation allows rapid ecotype formation"

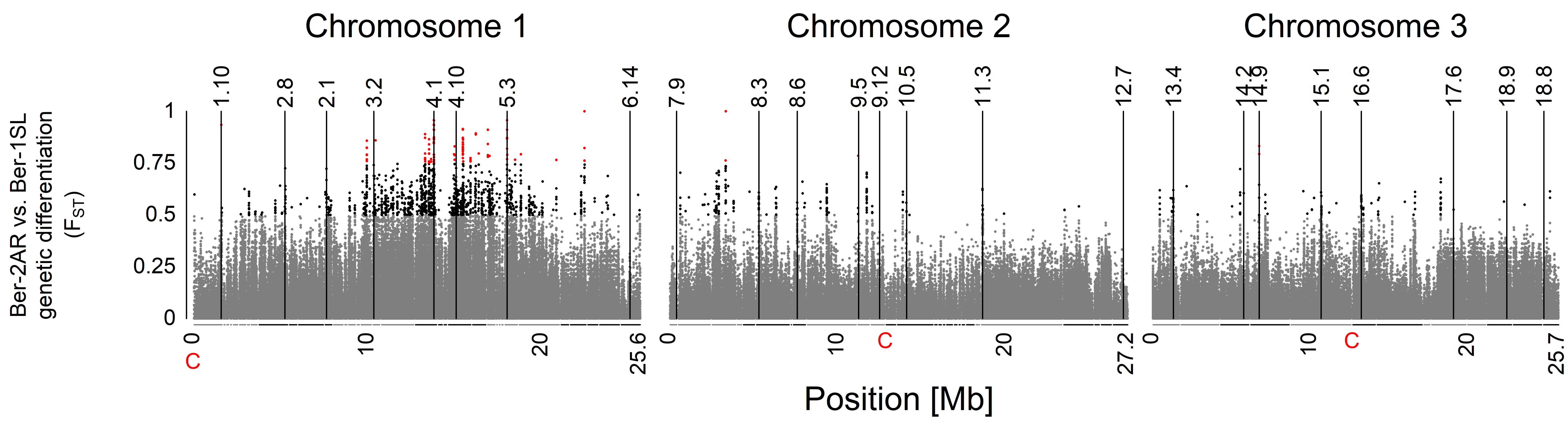


**Supplementary Figure 21 Location of the designed genetic markers across the genome of *Clunio*.**

The Manhattan plot shows the genetic differentiation ($F_{ST}$) between the populations Ber-2AR and Ber-1SL. The $F_{ST}$ values are color-coded in grey ($F_{ST}$ < 0.5), black (0.5 <= $F_{ST}$ < 0.75) and red ($F_{ST}$ <= 0.75). The genetic markers are marked with black lines at their exact location on the chromosomes. The scaffolds of the genome are subdivided by colors to indicate approximately equally sized regions on the chromosomes.
