## Supplementary Figure 22 for "Polygenic adaptation from standing genetic variation allows rapid ecotype formation"

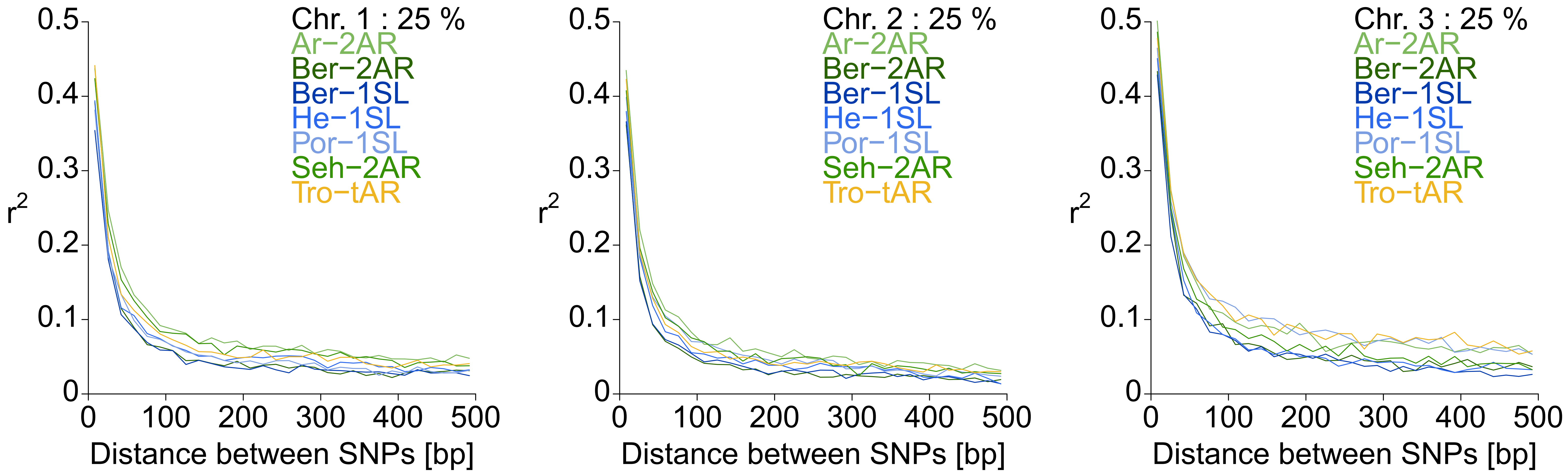


**Supplementary Figure 22 Linkage disequilibrium (LD) decay in the studied *C. marinus* populations.**

Populations were analyzed separately and the corresponding lines in each plot are color-coded. To make a pairwise LD calculation measured as $r^2$ feasible, the SNP set was randomly subsampled to 25%. The distance between the SNPs was limited to 500 bp for LD calculation. The resulting $r^2$ values were then averaged within 30 average-distance-classes.
