## Supplementary Table 1 for "Polygenic adaptation from standing genetic variation allows rapid ecotype formation"

**Supplementary Table 1 Wilcoxon rank sum test with continuity correction for significant differences in nucleotide diversity (π) between populations, based on the arithmetic mean of 200 kbp genomic windows.**

The lower-left part of the table shows the p-values and the upper-right part the corresponding significance levels: **** - 0-0.0001; *** - 0.0001-0.001; ** - 0.001-0.01; * - 0.01-0.05; ns - 0.05-1.

| | Por-1SL | He-1SL | Ber-1SL | Ber-2AR | Seh-2AR | Ar-2AR | Tro-tAR |
|---|---|---|---|---|---|---|---|
| Por-1SL | - | ns | ns | ns | **** | **** | **** |
| He-1SL | 0.06925 | - | ** | ns | **** | **** | **** |
| Ber-1SL | 0.3596 | 0.006378 | - | ns | **** | **** | **** |
| Ber-2AR | 0.619 | 0.1703 | 0.1576 | - | **** | **** | **** |
| Seh-2AR | 5,63E-10 | 4,87E-06 | 1,74E-13 | 1,93E-09 | - | *** | ns |
| Ar-2AR | 2.2E-16 | 1,74E-15 | 2.2E-16 | 2.2E-16 | 0.0001517 | - | * |
| Tro-tAR | 5,70E-15 | 5,68E-10 | 2.2E-16 | 1,27E-14 | 0.06954 | 0.03893 | - |
