## Supplementary Table 2 for "Polygenic adaptation from standing genetic variation allows rapid ecotype formation"

**Supplementary Table 2 SnpEff analysis for selected and all variants.**

SnpEff was run on the selected BayPass variants in association with the ecotype and all variants. For each analytical group (Region, Impact, Function, Variant) the total numbers for each subgroup and the fractions in respect to the entire analytical group are given. Additionally, the p-values for significant deviation between the selected and entire genome variants were calculated using Fisher's exact test.

| | Ecotype-associated variants (n=4,741) | | All variants (n=948,128) | | |
|---|---|---|---|---|---|
| | **n** | **%** | **n** | **%** | **p-value** |
| **Region** | | | | | |
| Exon | 1061 | 19.06 | 239000 | 20.69 | 0.002538563 |
| Intron | 1821 | 32.71 | 389723 | 33.74 | 0.1053189 |
| UTR | 418 | 7.51 | 98365 | 8.52 | 0.0069941 |
| Intergenic | 2209 | 39.68 | 413274 | 35.78 | 0.000000001811437 |
| Other | 58 | 1.04 | 14663 | 1.27 | 0.1490632 |
| **Impact** | | | | | |
| High | 31 | 0.56 | 10744 | 0.93 | 0.00251666 |
| Low | 612 | 10.99 | 141746 | 12.27 | 0.003404886 |
| Moderate | 479 | 8.6 | 101698 | 8.8 | 0.6185761 |
| Modifier | 4445 | 79.85 | 900837 | 77.99 | 0.0008337278 |
| **Function** | | | | | |
| Missense | 465 | 45.32 | 98560 | 43.06 | 0.1462238 |
| Nonsense | 2 | 0.19 | 1670 | 0.73 | 0.04045629 |
| Silent | 559 | 54.48 | 128647 | 56.21 | 0.2698506 |
| **Variant** | | | | | |
| SNP | 3890 | 82.05 | 792032 | 83.54 | 0.006363973 |
| InDel | 851 | 17.95 | 156096 | 16.46 | 0.006363973 |
