## Supplementary Table 4 for "Polygenic adaptation from standing genetic variation allows rapid ecotype formation"

**Supplementary Table 4 Sampling sites and sampling campaigns for the five newly established laboratory strains in this study.**

On sampling dates in squared brackets egg clutches for setting up laboratory cultures were collected from copulating pairs on the water surface. Samples from underlined dates were used for the genomic analyses of the wild populations.

| Sampling Site | Coordinates | | Ecotype | Laboratory Strain | Field Samples | Founding egg clutches |
|---|---|---|---|---|---|---|
| Sehlendorf (Germany) | 54° 18' 29" N | 10° 43' 30" E | Baltic | Seh-2AR | [05/2017], [06/2017] | 69 |
| Ar (Gotland, Sweden) | 57° 55' 06" N | 18° 56' 19" E | Baltic | Ar-2AR | [08/2018] | 36 |
| Kviturdvikpollen (Norway) | 60° 15' 58" N | 05° 14' 52" E | Baltic | Ber-2AR | 06/2017, [08/2018] | 40 |
| Kviturdvikpollen (Norway) | 60° 15' 58" N | 05° 14' 52" E | Atlantic | Ber-1SL | 06/2017, [08/2018] | 60 |
| Tromsø (Norway) | 69° 41' 04" N | 18° 53' 59" E | Arctic | Tro-tAR | [06/2018], [07/2018] | 47 |
