## Supplementary Table 5 for "Polygenic adaptation from standing genetic variation allows rapid ecotype formation"

**Supplementary Table 5 Multiplex PCR primer pairs for QTL mapping between the sympatric populations Ber-1SL and Ber-2AR.**

All primers were designed and tested by Kerstin Schaefer.

| Chromosome | Scaf old | Primer name | Sequence 5′-3′ | Product length |
|---|---|---|---|---|
| 1 | 8 | 1.10_f | TTCCTGGAAATACTGCAGCAA | 278 bp |
| | | 1.10_r | TCAACGAGATTTTCCGGTTTCA | |
| | 59 | 2.8_f | TGTCTTGCGAGGTTCTCTGTT | 450 bp |
| | | 2.8_r | CGCAATTTCAAGCTCTGATTGC | |
| | 6 | 2.1_f | ACACGAGATGAAACTCATGCG | 909 bp |
| | | 2.1_r | CCAAGGTCTACGAAAAAGCAGC | |
| | 53 | 3.2_f | CACGGTTAACATGCTTGAAGGA | 484 bp |
| | | 3.2_r | TCAAGTTCCAGCGCATTCGAT | |
| | 18 | 4.1_f | TGTGACTAACGGCAGGAAAGA | 782 bp |
| | | 4.1_r | ACAAAACGATTGGAAGCGACG | |
| | 43 | 4.10_f | CCCCGCAAATTTTCTTCGGAA | 537 bp |
| | | 4.10_r | ATCGGCGGTTGTTTTTGGTTT | |
| | 42 | 5.3_f | GGAATCGCACGAAGCTCATTT | 596 bp |
| | | 5.3_r | AACGCCACTAACACTCCAAAT | |
| | 61 | 6.14_f | AGCGACACAGATGAAAGACGA | 698 bp |
| | | 6.14_r | TGCGGCAATTTGTTTTAATGTCC | |
| 2 | 20 | 7.9_f | ACCTCAATATGCACGTCCAGT | 543 bp |
| | | 7.9_r | TCTGATATGGTCGCAAGTGGT | |
| | 56 | 8.3_f | CCCCAGCGCTTTTCACATTT | 369 bp |
| | | 8.3_r | TTTCCTCGGTTCTTCGTCCC | |
| | 21 | 8.6_f | TTTCACTGCCTTGTACACCCC | 451 bp |
| | | 8.6_r | ATGATTCGATGGGACGACTCG | |
| | 4 | 9.5_f | AGCTCCACTATTCCAAACAGCT | 727 bp |
| | | 9.5_r | GCCATCAGCCTTGAGAAATGC | |
| | 40 | 9.12_f | TGATCAACCAGTACAAGGTGGA | 424 bp |
| | | 9.12_r | GTAAGGAAGCATTAGCAGCGC | |
| | 50 | 10.5_f | CGCTACTTTTGCCCATCAACA | 478 bp |
| | | 10.5_r | TGTCGCCATCACTTTGGAATG | |
| | 47C | 11.3_f | CATTGCTTGCGATTTGCTTTGA | 455 bp |
| | | 11.3_r | GGAATCTCGTGCATCGACGAT | |
| | 47F | 12.7_f | GCCTTGCAATAGCGCATACG | 849 bp |
| | | 12.7_r | TAGGCTCTGGGGTATGGCTT | |
| 3 | 10 | 13.4_f | GTTCCTTGCCACTCATTTGCT | 450 bp |
| | | 13.4_r | TGTCGCCAAACAGGATTTGTG | |
| | 41A | 14.2_f | TGGCAAATCAAGGACCAGACA | 740 bp |
| | | 14.2_r | TCGACTTGAGGCATTATTGGGA | |
| | 41B | 14.9_f | TCTTGCGGCAAGATCCACTTT | 606 bp |
| | | 14.9_r | TAAACGATTCAGTTCGCACGC | |
| | 48 | 15.1_f | CTCGATTCAGACCGTCCGATT | 639 bp |
| | | 15.1_r | GGCGTGACCTTCATGAAGAGA | |
| | 51 | 16.6_f | GCTGAATAAGGCATTGCTGGG | 492 bp |
| | | 16.6_r | GCAGAGTAGTGCGAGTGGAAT | |
| | 45B | 17.6_f | TCTCACTTTTCACGACGCTTTT | 308 bp |
| | | 17.6_r | TAGAAGCTGTTCCGCCTCAAG | |
| | 5 | 18.9_f | CAACGTGAGACGAACGAACAG | 543 bp |
| | | 18.9_r | CGTGGCCTTGTTTCAACACAA | |
| | 3 | 18.8_f | CCACTGAAACCATCGGGACA | 227 bp |
| | | 18.8_r | AGAATGTGTGACTGTTGCAAAGT | |
