## Supplementary Table 6 for "Polygenic adaptation from standing genetic variation allows rapid ecotype formation"

**Supplementary Table 6 Volume modifications for the REPLI-g Mini Kit QIAGEN 150025 whole genome amplification kit (QIAGEN).**

MM – Master Mix.

| Phases | template DNA | Buffer D1 | Buffer N1 | MM: Nuclease-free water | MM: REPLI-g Mini Reaction Buffer | MM: REPLI-g Mini DNA Polymerase | MM: Adding total volume |
|---|---|---|---|---|---|---|---|
| Manufacturers' Volume (suitable for 15 reactions) | 2.5 µl | 2.5 µl | 5 µl | 10 µl | 29 µl | 1 µl | 40 µl |
| Edited Volume | 7.5 µl | 1 µl | 1.5 µl | 0 µl | 14.5 µl | 0.5 µl | 15 µl |
