## Supplementary Table 7 for "Polygenic adaptation from standing genetic variation allows rapid ecotype formation"

**Supplementary Table 7 The three covariates used for association analysis in BayPass.**

| Covariates | Ar-2AR | Seh-2AR | Ber-2AR | Ber-1SL | He-1SL | Por-1SL |
|---|---|---|---|---|---|---|
| Ecotype (binary) | 0 | 0 | 0 | 1 | 1 | 1 |
| Sea Surface Salinity (PSU) | 7 | 14.5 | 31.5 | 31.5 | 31.3 | 33.9 |
| Average Water Temperature 2020 (°C) | 7.5 | 9.1 | 8.9 | 8.9 | 10.1 | 11.8 |
