## Supplementary Note 1 for "Polygenic adaptation from standing genetic variation allows rapid ecotype formation"

**Supplementary Note 1: Oviposition behavior in the Baltic ecotype**

Because of the lack of tides in the Baltic Sea, the Baltic ecotype cannot rely on the tides to expose the larval substrates for egg deposition. Therefore, in contrast to the Atlantic and Arctic ecotypes, the Baltic ecotype does not oviposit on exposed larval substrates, but on the water surface, from where the eggs sink to the bottom of the sea. This change in oviposition preference is accompanied by a specific behavioral change, namely the bending of the female’s abdomen down through the water surface ^1^, which ensures that the egg masses are not caught in the water’s surface tension. Additionally, the Baltic ecotype’s egg jelly is reported to have slightly differing properties ^1^.

In our laboratory culture, the animals are kept in plastic boxes with a constant water level, so that larval substrates are never exposed. Under these conditions, most of the Baltic ecotype’s egg masses will be found submerged at the bottom of the culture box, as expected. Very rarely egg masses are deposited on the walls or the lid of the box. The Atlantic and Arctic ecotypes will – in the absence of exposed larval substrates – also oviposit on the water surface. However, as they lack the characteristic banding of the female abdomen during oviposition, their egg masses will be trapped in the water’s surface tension and cannot sink to the bottom of the culture box. Their egg masses will always float on the water surface, usually with the female still sticking to the egg jelly. These floating egg masses often form aggregations on the water surface.

We used these differences in where the egg clutches are found in our laboratory cultures to additionally confirm ecotype identity of our laboratory strains.

1	Endraß, U. Physiologische Anpassungen eines marinen Insekts. II. Die Eigenschaften von schwimmenden und absinkenden Eigelegen. *Marine Biology* **36**, 47-60 (1976).
